# Inferring Ecosystem Networks as Information Flows

**DOI:** 10.1101/2021.02.18.431917

**Authors:** Jie Li, Matteo Convertino

**Author notes:** Corresponding author: M. Convertino, 9 Chome, Kita 14, Nishi 9, Kita-ku, room 11-11, Graduate School of Information Science and Technology, Hokkaido University, Sapporo, Hokkaido, Japan, 060-0814.

## Abstract

The detection of causal interactions is of great importance when inferring complex ecosystem functional and structural networks for basic and applied research. Convergent cross mapping (CCM) based on nonlinear state-space reconstruction made substantial progress about network inference by measuring how well historical values of one variable can reliably estimate states of other variables. Here we investigate the ability of a developed Optimal Information Flow (OIF) ecosystem model to infer bidirectional causality and compare that to CCM. Results from synthetic datasets generated by a simple predator-prey model, data of a real-world sardine-anchovy-temperature system and of a multispecies fish ecosystem highlight that the proposed OIF performs better than CCM to predict population and community patterns. Specifically, OIF provides a larger gradient of inferred interactions, higher point-value accuracy and smaller fluctuations of interactions and *α*-diversity including their characteristic time delays. We propose an optimal threshold on inferred interactions that maximize accuracy in predicting fluctuations of effective *α*-diversity, defined as the count of model-inferred interacting species. Overall OIF outperforms all other models in assessing predictive causality (also in terms of computational complexity) due to the explicit consideration of synchronization, divergence and diversity of events that define model sensitivity, uncertainty and complexity. Thus, OIF offers a broad ecological information by extracting predictive causal networks of complex ecosystems from time-series data in the space-time continuum. The accurate inference of species interactions at any biological scale of organization is highly valuable because it allows to predict biodiversity changes, for instance as a function of climate and other anthropogenic stressors. This has practical implications for defining optimal ecosystem management and design, such as fish stock prioritization and delineation of marine protected areas based on derived collective multispecies assembly. OIF can be applied to any complex system and used for model evaluation and design where causality should be considered as non-linear predictability of diverse events of populations or communities.

---

*“But truth is ever incoherent”*

Astrid Recker

## 1 Introduction

### 1.1 Ecosystem Complexity and Predictability

The flourishing development of complexity science ^1;2^ has shed light on research questions and applications in many interdisciplinary fields, for instance, climate change^3;4;5^, epidemiology ^6;7^ and ecosystem sciences at multiple scales ^8;9;10^. In this burgeoning science, complex network models play a central role in the quantitative analysis, synthesis and design (including predictions) of ecosystems and their visual representation. This is because functional and structural networks – such as species interactions and ecological corridors – are the core elements of ecosystems defining species organization and ecosystem function. When inferring networks, causal inference^11^ is one of the fundamental steps for ecosystem reconstruction and graphical representation by assessing interactions or interdependencies dynamically or across a period of time – between biota, environment, and among those – that can be conceptualized as information flows in a general purview^12^. In a quantitative sense, network inference can performed via causality inference based on time series data defining the dynamics of ecosystem components. Causal inference also attracts much attention in some emerging disciplines such as big data science via machine learning since it brings a new set of tools and perspectives for some problems in these areas. However, this issue of causal inference is still an extremely challenging problem due to the intrinsic lack of knowledge or observability of the “true” reality of a system especially for highly complex non-linear systems driven by nonlinear environmental forcing. Certainly the objective of causal inference is defining unknowns; however robust model validation must be performed. In order to make causal inference practical and achievable, causality is often replaced with predictability as it is articulated in this paper. A plethora of conceptual approaches, frameworks and algorithmic tools including but not limited to Pearson’s Correlation Coefficient (PCC)^13;14^, Bayesian Networks (BNs) and Dynamic Bayesian Networks (DBNs) ^15;16;17;18;19^, Neural Networks, Graphical Gaussian Models (GGMs) ^20;21^, Wiener-Granger causality (GC) model ^22^, Structural Equation Modeling (SEM) ^23;24;25;26;27;28^, Convergent Cross Mapping (CCM)^29^ and information-theoretic models ^30;31;32;33^ for instance, to tackle causal interactions and infer complex networks in terms of correlation, predictability and probability have been well established; however, most tools are solely tested on low-dimensional systems and some are even untested on ecosystems at different levels of complexity or simulated ones. The vast majority of these models in ecosystem science (with the exception of CCM and few others such as PCMCI^34^) consider only the inferred causality between species pairs one at at time without the simultaneous consideration of all species pairs for each species that is shaping ecosystem collective behavior mediated by environmental dynamics. The ensemble of all species causations is representable as nonlinear dynamical network over the space-time-environmental domain considered. It is therefore valuable for science to seek for robust models and explore novel methods to identify and quantify the pattern-oriented causality between variables (such as species) and how this causality is predictive of target complex system patterns.

### 1.2 Causal Interaction Inferential Models

#### 1.2.1 From Granger to Convergent Cross Mapping

For quite a long time, correlation has been considered as a heuristic of causal relationship between variables even though George Berkeley ^35^ always suggested that correlation did not necessarily or sufficiently imply causation. Especially for ubiquitous nonlinear dynamics, applying linear correlation to infer causation is cursory and risky. Statements about causation and correlation actually do not have much to do with each other, particularly when there is no a-priori knowledge of the studied ecosystem processes. The conceptualization and identification of “causality” was originally introduced by Wiener^36^ who propounded that “causality” between two variables can be identified by measuring how well one variable facilitates the predictability of the other. In this broader view causality was already conceptualized correctly as predictability.

In 1969, Granger formalized Wiener’s idea^36^ in terms of autoregression and established the framework of Wiener-Granger causality (GC) model ^22^ that after lead to the Nobel prize in economics. Since then, GC approach has become a frequently used advance for causation and useful to infer causal interactions between strongly coupled variables. According to the concept of GC model, a variable is said to “GC cause” a second variable if knowledge of the current value of the first variable helps in predicting that of the second variable. This notion of causality was substantially based on the predictability of time series, although strictly speaking Granger causality is about conditional independence of variables rather than predictability. The key requirement of GC model is separability that is a feature of purely linear and stochastic systems ^22^, and provides a way to understand the system as sum of components rather than as a whole non-linear entity composed by multiple components difficult to separate. Separability means that the second variable can be independently and uniquely forecasted by the first variable; an assumption that reflects how studied systems are interpreted as linear systems and that is certainly not the case of real complex systems. Additionally, states in the past of some variables in dynamical systems can be inherited through time, which means that the behavior of dynamical systems has memory. Yet, both cause and effect are embedded in a non-separable higher dimension trajectory. Space-time separability therefore becomes extremely hard to satisfy in systems that can be described as complex networks where each node (variable) influences several nodes or even all nodes in the entire system simultaneously, resulting in a non-random propagation of information through the network. In this sense ecosystems can be thought that information machines where separability is only possible by fixing thresholds of significance for the patterns to investigate. As a consequence, GC model might be problematic while using in nonlinear dynamical systems with deterministic settings and weak to moderate interaction. In attempt to solve the causality inference problem in complex ecosystems, Sugihara et al. (2012) ^29^ developed the Convergent Cross Mapping (CCM) model ^29^, and successfully applied this model to a coupled non-linear mathematical predator-prey model and a real-world sardineanchovy-temperature ecosystem. Later on, Ushio et al. (2018) ^37^ applied CCM to a complex fish ecosystems with 15 species after removing seasonality from abundance data in order to assess “true” or biological interactions.

In dynamic systems two variables (X and Y for instance) are causally linked if they are generated by one system and share a common attractor manifold. It implies that each variable can be used to recover (predict) the other one. CCM is the method capable of quantifying this kind of correspondence between two variables. CCM does so by measuring the extent to which the states of one variable (considering values rather than probability distributions) can be reliably estimated by the other one with time lags. In practice, CCM take values of variables X and Y, a time lag embedding is derived from the time series of Y, and the ability to estimate the states of variable X from the time lag embedding quantifies how much signature of X is encoded in the time series of Y. This principle was termed as “Cross Mapping”, and it was suggested that the causal effect of X on Y is determined by how well Y “cross maps” X. Sugihara et al. (2012) ^29^ noted that CCM had drawbacks, although some of these are disputable. For instance for the phenomenon of “generalized synchrony” as a result of exceptionally strong unidirectional causation (X strongly “cross maps” Y, but Y does not causes X). In such a case, both directions (X “cross maps” Y, Y “cross maps” X) of the causal relationship can be observed from CCM’s results, resulting in a “misleading” bidirectional causality ^38^. This was perceived as a limitation of CCM in distinguishing between bidirectional causality and strong unidirectional causality because of the synchrony. Misleading is however not a correct definition since we believe any variable has always non-zero interdependencies due to unaccounted factors and chance that interactions may appear at least once in the ecosystem considered. Yet, asymmetrical interdependence is a norm rather than a numerical artifact. Another key property, and potentially a drawback, of CCM is convergence that is stable predictability (rho) after a critical library size defining the minimum information for reliable inference. However, datasets are not always long enough especially for real-world applications. Yet, convergence might be limited by the finite size of time series data. Lastly, CCM suffers from the high computational complexity in terms of model parameters and computation speed. Despite these drawbacks CCM is used in this study as a benchmark for evaluating our proposed model.

### 1.2.2 From Information Flow to Transfer Entropy

Most natural and artificial systems composed of a large number of interacting elements can be represented as networks. Networks, functional and structural, are the backbone of ecosystems. In order to untangle such networks, the primary mission is to identify and quantify causations between elements and then to infer the networks for analysis and visualization. These networks are information fluxes representing ecosystems via non-linear dynamic interactions. Ecosystems can be identified in terms of network topology as collective dynamics and as a function of macroecological indicators as entropy/energy states. Variables in information theory including Shannon entropy, Mutual Information (MI) and Transfer Entropy (TE) have been recently used in complex network science to characterize ecosystems at different scales ^39^. Herein, TE coined by Thomas Schreiber^40^ is an information-theoretic quantity measuring the asymmetric bidirectional information transfer (vs. information flow as in Lizier et al. (2010) ^41^ when conditional entropies are used to exclude indirect pairs of species whose interactions is of second order importance) between two variables ^42^. In a conceptual and practical view, besides GC and CCM, TE can be an appropriate candidate to infer the causality between interacting elements in complex ecosystems.

As mentioned above, GC model may be problematic in complex systems due to highly nonlinear dynamics, and CCM may not be suitable for distinguishing well bidirectional or strong unidirectional interactions, the requirement convergence, the lack of consideration of probability distribution functions (pdfs), and numerical sensitivity due to high computational complexity. By contrast, TE, as a non-parametric, model-free information-theoretic variable defined from nonlinear dynamics of Markov chain process (mappable as stochastic pdf propagation equivalently), it provides a directed measure to detect asymmetric dynamical information transfer between two time-varying variables. TE is particularly convenient because, on the contrary of other models such as CCM, is a “first principle” variable defined without assuming any particular functional/process or numerical model to identify the interactions in studied systems ^43^. For this reason, TE is the elementary block of complex systems represented as information processing machines where uncertainty or information shapes systems’ collective behavior driven by non-linear convolution of intrinsic system’s properties and external noise (e.g. biology and environment, respectively). Thus, TE has been considered as an important and powerful tool to analyze causal relationships in nonlinear complex systems^44^ and numerical methods are just used to calculate TE from its analytical form (such as the choice of pdf binning and entropy discretization). It is worth noting that for Gaussian random processes TE is equivalent to Granger causality ^45^ but these are rare processes in nature. The vast majority of natural process are multimodal and non-Gaussian.

Although, in its basic formulation, TE has already been widely used for causality inference, general principles, unified frameworks, and models, and further developments based on TE are still lacking. More importantly no work hitherto has been done to give systematic validations for TE-based causality inference models with mathematically synthetic data, as well as real-world ecosystems, to elucidate how TE behaves dependent on dynamics and complexity. Razak et al. (2014) ^46^ made progress on this issue by using classical and amended Ising models which are mathematical models of ferromagnetism in statistical mechanics; Duan et al., (2013) ^47^ provided a theoretical and experimental systemic validation of a TE-based model; and finally Runge et al. (2018) ^48^ explored TE and other models with synthetic data. However, these studies are applied to complex systems with a limited number of variables or whose dynamics is well defined; yet, they did not validate the model for realistic ecosystems in its full complexity, driven also by data fallacies, as seen in nature. Therefore, specific applications of TE-based models lack of a rigorous performance assessment that thus remains elusive. On one side low complexity well-known ecosystems can validate the inferred pairwise interactions, while highly complex ecosystems can validate the whole systemic interaction network predictability on some patterns such as biodiversity indicators over time. The former problem deals more with accurate causality between pairs, while the latter deals more with ecosystem predictability.

### 1.3 Optimal Information Flow and Predictability

In this study, to overcome limitations of CCM and TE inference models estimating species pairs independently of each other, as well as to revise and validate “causality” in a predictive sense, we propose the Optimal Information Flow (OIF) model based on previously developed models by our group ^49;39^. Specifically, the proposed OIF model involves four main steps: (i) MI-based optimal time delay assessment that maximizes the predictability of variables and focus on extreme or rare interactions^39^; (ii) TE computation considering data probabilistic dynamics; (iii) coupled maximization of uncertainty reduction and removal of indirect links; and (iv) selection of optimal TE threshold for predicting selected patterns (e.g. *α*-diversity). The optimal threshold on TE is not necessarily within the scale-free maximum uncertainty reduction range (describing ecosystem collective dynamics) because predicted patterns define the patter-specific salient TE. Technically, in this paper we offer a different estimation of TE than Li and Convertino (2019) ^39^ (based on JIDT Kernel estimator suitable for powerlaw distributed data^42^), and we explore all TEs (without TE thresholding) with the aim of capturing all differences between TEs and CCM interaction matrices and their ability to predict fluctuations in macroecological patterns considering connected species a posteriori. In addition, OIF is improved with respect to Li and Convertino (2019) ^39^ by considering its extension over time to reconstruct dynamical information networks, and the time-dependent Markov order (self-memory) of each species. Note that in this paper we consider all inferred TEs without any redundancy check and removal of indirect interactions as in Servadio and Convertino, (2018) ^49^ because we wish to characterize the full interaction matrix without any assumptions on biological interactions or methodological criteria of subordinate interactions. This is particularly important when no knowledge is available a priori about biological interactions, whether the biomarker used (e.g. abundance) is reflective of the interactions of interest (physical, biomass conversion, hormonal interactions, etc.), and when indirect weak interactions are quite important (and this is quite common for small organisms, e.g. microbes).

The performance of OIF is assessed by applying it to three prototypical case studies including mathematical deterministic and real-world ecosystems. One is a biologically inspired mathematical model that can generate synthetic two-coupled time-series variables describing dynamics similar to predator-prey dynamics^29^. Two parameters (*β_xy_* and *β_yx_*) in the equations underlying the model are describing the strength of true interdependence between two simulated variables and they can be free varied. Other two case studies are real-world ecosystems: the case of externally forced poorly coupled species (sardine-anchovy-temperature system) ^29^ and the one of highly complex interacting species (fish community in Maizuru bay)^37^ (Fig. 1). The well-documented CCM method is also used for these three cases and the results from CCM, despite its known drawbacks of convergence, strong asymmetrical causality miscalculation, and computational complexity, were somewhat considered as benchmark interactions due to the lack of other estimates.

**Figure 1.**
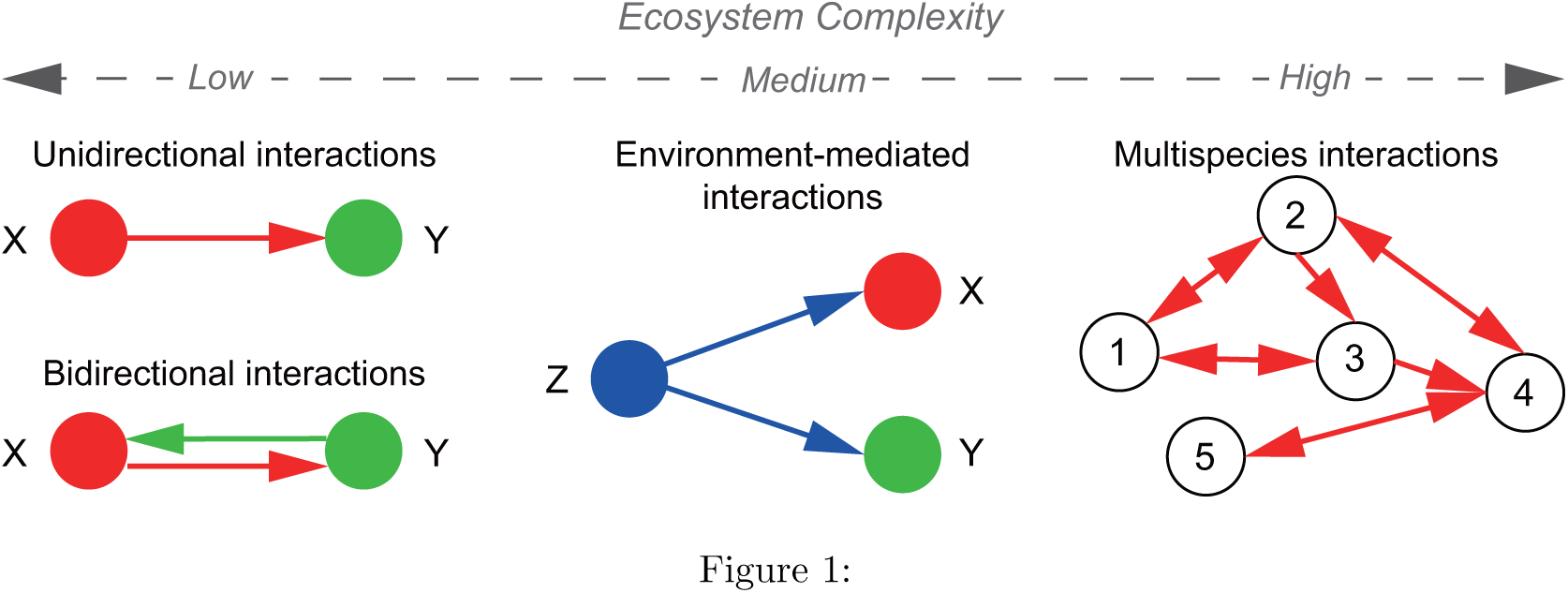
Studied ecosystem complexity. Epitomes of increasing ecosystem complexity are shown from left to right where nodes are representing variables (e.g. species or other socio-environmental features). Case 1 shows two basic cases: unidirectional and bidirectional interactions where true interaction strength is known because embedded into a mathematical model. Case 2 is about environment-mediated interactions with no knowledge of “true” interactions. Case 3 is a multispecies ecosystem with multiple bidirectional interactions with no knowledge of “true” interactions.

OIF is not perceived as a competitor with other already published models, including GC and CCM, but rather it aims at providing an alternative and hopefully more precise assessment to predictive inference in cases not completely covered by previous models. Theoretically, leaving aside systematic data issues, OIF is expected to give a better performance than other models in interdependence assessment owing to the aforementioned fine properties of TE for nonlinear dynamics. Besides, given the relationship between entropy and diversity ^50^ (specifically Shannon and Transfer entropy and *α*- and *β*-diversity), OIF provides a potential advantage to predict the information about macroecological indicators of ecosystems. In consideration of these features, TE causality is proposed as *non-linear predictability* of both population fluctuations of species (in this case abundance fluctuations) and community macroecological indicators, simultaneously.

The fundamental principle about OIF is anchored into the idea of systemic uncertainty reduction (leading to maximum predictive accuracy) that is by itself a form of quantitative validation considering aleatoric uncertainty in species variables. Certainly systematic uncertainty (of models and algorithms) and biological ignorance are two other important elements to consider for a complete validation, and we tried to address those to a certain extent but further studies are required. It should be kept in mind that Interactions are always specific to a target outcome, e.g. biomass variation, that is likely reflected by the species biomarker as model inputs. For this reason we emphasize that the inferred interactions are generally *predictive causal relationships* which maximize predictive accuracy of multivariate time series and not “absolute” species interactions (for an interesting semantics discussion on species interactions see Nakazawa (2020) ^51^).

## 2 Results

### 2.1 Two Species Unidirectional Coupling Ecosystem

This bio-inspired ecosystem S(*β_xy_* = 0, *β_yx_*) describing the unidirectional coupling is run for 1000 time steps for reaching stationarity, generating a set of 1000 points long time-series dependent on *β_yx_*. This means that species X has an increasing effect on Y with the increase of *β_yx_*, but Y has no effect on X. Both CCM and the proposed OIF model are separately used to quantify the potential causality between species X and Y. The inferred causality dependent on *β_yx_* only (as a physical interaction) is shown in Fig. 2A. *β*, *ρ* and TE have different units; specifically *β* and *ρ* are dimensionless while TE is measured in bits or nats (a logarithmic unit of information or entropy). Therefore, any comparison is done considering gradients of change when these variables vary together rather than making comparisons between absolute values which are meaningless. Figure 2A shows that under the condition of *β_xy_* =0, results of “Y to X” (i.e. the estimated effect on Y on X) is close to 0 for the OIF model (*T E_Y →X_* (*β_yx_*)) that precisely describe the no-effect of Y on X. “X to Y” (*T E_X→Y_* (*β_yx_*)) well tracks the increasing strength of the effect of X on Y for increasing values of the physical interaction *β_yx_* embedded into the mathematical model. However, considering results of the CCM model, “Y to X” (*ρ_Y →X_* (*β_yx_*)) presents non obvious (and likely wrong) non-zero values with higher fluctuations compared to *T E_Y →X_* (*β_yx_*) especially for lower values of *β_yx_*. This erroneous estimates of CCM is likely related to the need of CCM for convergence. For CCM, “X to Y” ((*ρ_X→Y_* (*β_yx_*))) shows an increasing trend for increasing values of *β_yx_* and decreasing when *β_yx_* is greater than *∼*0.5 non-trivially. In consideration of these results for the unidirectional coupling ecosystem, the OIF model performs better over CCM in terms of unidirectional causality inference.

**Figure 2.**
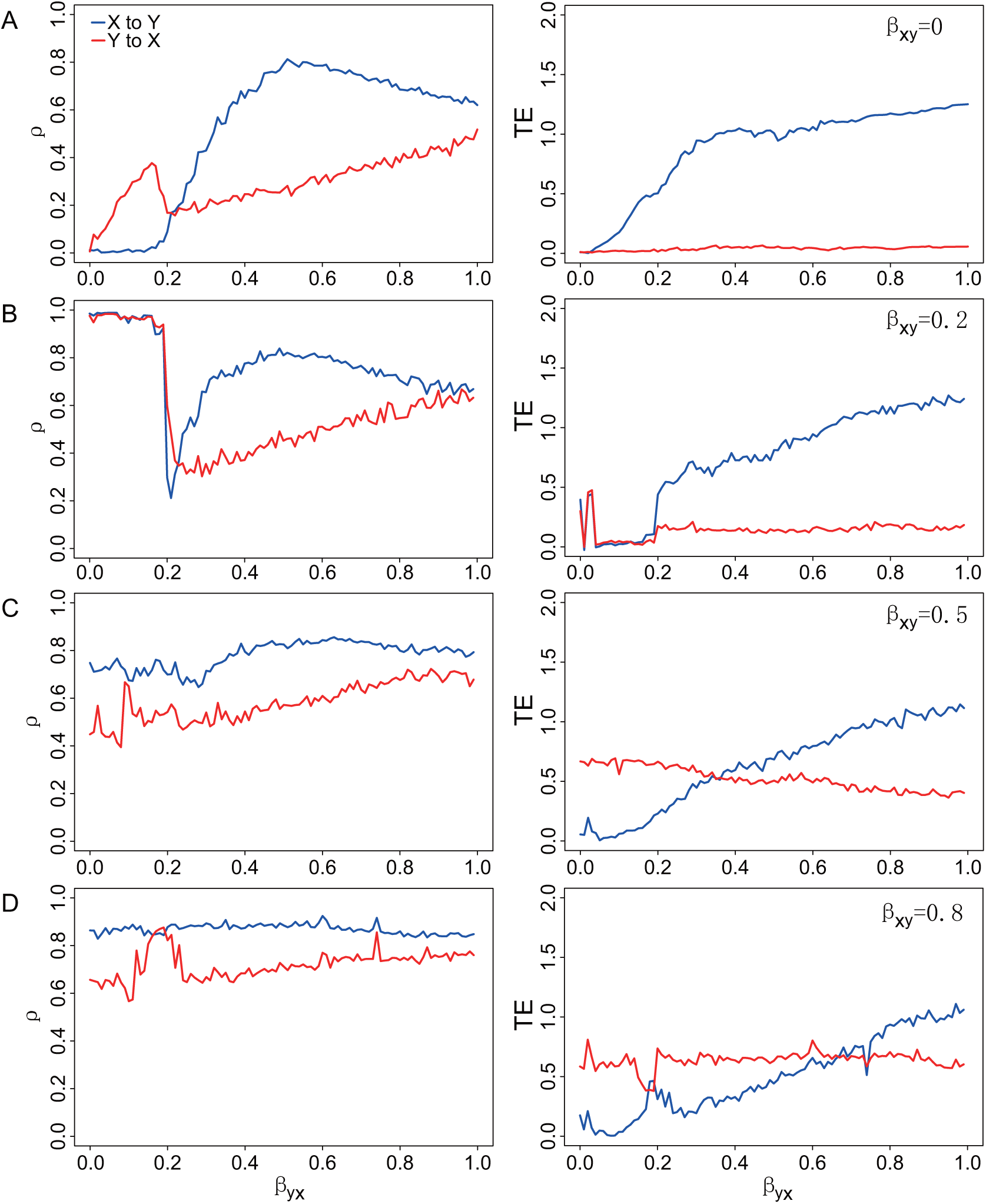
Inferred predictable causality via CCM and TE for embedded true causality. CCM correlation coefficient (*ρ*, left plots) and Transfer Entropy (TE, right plots) are shown for the bio-inspired mathematical model in Eq. 5.1 representing bidirectional interactions. The mathematical model indicated as S(*β_xy_*, *β_yx_*) is simplified as a univariate function because *β_xy_* is fixed while *β_yx_* is free and varying within the range [0, 1]. *β_xy_* and *β_yx_* are establishing true causality while *ρ* and TE are indicators of predictable causality. Y’s causal effects on X is theoretically fixed as a stable value corresponding to each *β_xy_*. The greater *β_xy_* the stronger Y affects X (estimated by *ρ_yx_* and *T E_yx_* in red lines). (A) *β_xy_* = 0 means that Y does not affect X and then X dynamics is only related to stochastic dynamics due to birth-death process as in the model (Eq. 2.1). X’s effects on Y depends on the value of *β_yx_*, theoretically leading to increasing functions *ρ_xy_* and *T E_xy_* (blue lines) when *β_yx_* increases; (B) *β_xy_* = 0.2; (C) *β_xy_* = 0.5; and (D) *β_xy_* = 0.8.

### 2.2 Two Species Bidirectional Coupling Ecosystem

In this case, the effect between two species is bidirectional. Species X has an effect on species Y and vice versa. The univariate dynamical systems S(0.2/0.5/0.8, *β_yx_*) are run for 1000 time steps under the same conditions determined by *β_xy_*. Certainly this situation is fictional since in real ecosystems the interaction strength is changing when other interacting species change their interactions.Thus, keeping one interaction fixed around one value is a strong unrealistic simplification (analogous of one-factor at-a-time sensitivity analyses) but it is a toy model that allows to verify the power of network inference models. These models generate three sets of 1000 points long time-series dependent of *β_yx_* for each fixed *β_xy_*. OIF and CCM are used to infer “causality” between X and Y – in the form of *ρ* and TE – and compare that against the real embedded interaction *β_yx_* and *β_xy_* shown in Fig. 2BCD. Considering all results of Figure 2 corresponding to fixed *β_xy_* s, the correlation coefficient *ρ* yielded from CCM and TE from OIF are both able to track the strength of causal trajectories. However, TE seems to perform better in term of ability to infer fine-scale changes in interactions. In particular, considering Fig. 2D (right plot), higher *T E_yx_* higher for low *β_yx_* makes sense because *β_xy_ > β_yx_* that means Y has a larger influence on X than vice versa and then Y is able to predict X. Additionally, TE does not suffer of convergence problems; specifically, considering Fig. 2A (left plot), higher *ρ* for small *β_yx_* is not sensical and that is likely related to convergence problems of CCM.

Considering all results of Figure 2 corresponding to fixed *β_xy_* s, the correlation coefficient *ρ* yielded from CCM and TE from OIF are both able to track the strength of causal trajectories. Ideally, the causality from Y to X is a constant since *β_xy_* is a fixed value for each case. In this figure, the red curve in the right panel representing the OIF-inferred (TE-based) causality from Y to X is higher for greater *β_xy_* s, while red curves representing CCM-inferred (*ρ*) causality in the left panel present higher fluctuations especially for lower *β_yx_*. For the causality from X to Y determined by *β_yx_* in the mathematical model, theoretically speaking, the causality from X to Y should monotonously grow when *β_yx_* increases from 0 to 1. In Figure. 2, blue curves in the right panel representing the OIF-inferred (TE-based) causality from X to Y present monotonously increasing features as a whole with the increasing *β_yx_*, while those from CCM model (*ρ*) do not and show considerable fluctuations.Therefore, OIF outperforms CCM in terms of the ability to infer the fine-scale changes in causality. In particular, considering Fig. 2D (right plot), higher *T E_yx_* higher for low *β_yx_* makes sense because *β_xy_ > β_yx_* that means Y has a larger influence on X than vice versa and then Y is able to predict X. Additionally, TE does not suffer of convergence problems; specifically, considering Fig. 2A (left plot), higher *ρ* for small *β_yx_* is not sensical and that is likely related to convergence problems of CCM.

Additionally, *ρ_Y →X_* (*β_yx_*) shows higher fluctuations on average especially for the condition of lower *β_yx_*s compared to *T E_Y →X_* (*β_yx_*). When considering the effect of X on Y that is a function of *β_yx_* for CCM, *ρ_X→Y_* reaches an extreme value at around *β_yx_* = 0.5 and then declines for larger values of *β_yx_*. This is not consistent with the expected effect of X on Y that should be proportional to *β_yx_* embedded into the mathematical model. The ability of *ρ* to reflect the proportional relationship between the effect of X on Y (manifested by *β_yx_*) vanishes for high *β_xy_* s due to unexpected and somewhat inconspicuous changes in *ρ_X→Y_* for larger *β_yx_*. In simple words, the expected increasing trend of *ρ* is lost for larger *β_xy_* that is counterintuitive. On the other side, *T E_X→Y_* (*β_yx_*) invariably maintains an increasing trend for increasing values of *β_xy_*. OIF is also performing better than CCM when predicting higher average values of *T E_Y →X_* for increasing values of *β_xy_* (red curves in Fig. 2ABCD, right plots) as expected by the fixed effect in the mathematical model of Y on X. These results suggest that when compared to *ρ* of CCM, TE can track well the causal interactions over *β_yx_* with higher performance and without considering the convergence requirement of CCM. CCM needs to consider the length of time series that makes *ρ_X→Y_* (*β_yx_*) convergent to a stable value, but uncertain for large differences in time-series length of (X,Y) and sensitive to short time series.

In more realistic settings for real ecosystems (and in analogy to global sensitivity analyses) when *β_xy_* and *β_yx_* are both considered as arguments of the two-variable (X,Y) bio-inspired model, the simulated ecosystem becomes a truly bivariate system, yet yielding complexity but more interest into the causality inference (Fig. 3). The dynamical system S(*β_xy_*, *β_yx_*) was generated for 800 time steps under the same conditions mentioned above. We generated the datasets that allowed us to study linear and non-linear predictability indicators for inferring the embedded physical interactions. Specifically, we measure undirected linear correlation coefficient *corr_X_*_;_*_Y_* (*β_xy_, β_yx_*), non-linear undirected mutual information *MI_X_*_;_*_Y_* (*β_xy_, β_yx_*), directed non-linear correlation coefficient *ρ_X→Y_* (*β_xy_, β_yx_*) and *ρ_Y →X_* (*β_xy_, β_yx_*), and non-linear directed transfer entropy *T E_X→Y_* (*β_xy_, β_yx_*) and *T E_Y →X_* (*β_xy_, β_yx_*) as shown in Fig. 3. These 2D phase-space maps in Fig. 3 show strikingly similar patterns for classical linear correlation coefficients, MI, *ρ* of CCM and TE of OIF which underline the fact that all methods are able to infer the interdependence patterns of interacting variables explicitly defined by *β_xy_* and *β_yx_*. The color of phase-space maps is proportional to the inferred interaction between X and Y when the mutual physical interactions are varying according to the mathematical model in Eq. 5.1. In Fig. 3, even though phase-space maps of undirected *corr_X_*_;_*_Y_* (*β_xy_, β_yx_*) and *MI_X_*_;_*_Y_* (*β_xy_, β_yx_*) present similar patterns (in value organization and not value range) to those of directed *ρ* and *T E*, neither *corr_X_*_;_*_Y_* ((*β_xy_, β_yx_*) and *MI_X_*_;_*_Y_* ((*β_xy_, β_yx_*) provide information about the direction of causality. As expected MI shows the opposite pattern of the average TE due to the fact that MI is the amount of shared information (or similarity) versus the amount of divergent information (divergence and asynchronicity) between X and Y.

**Figure 3.**
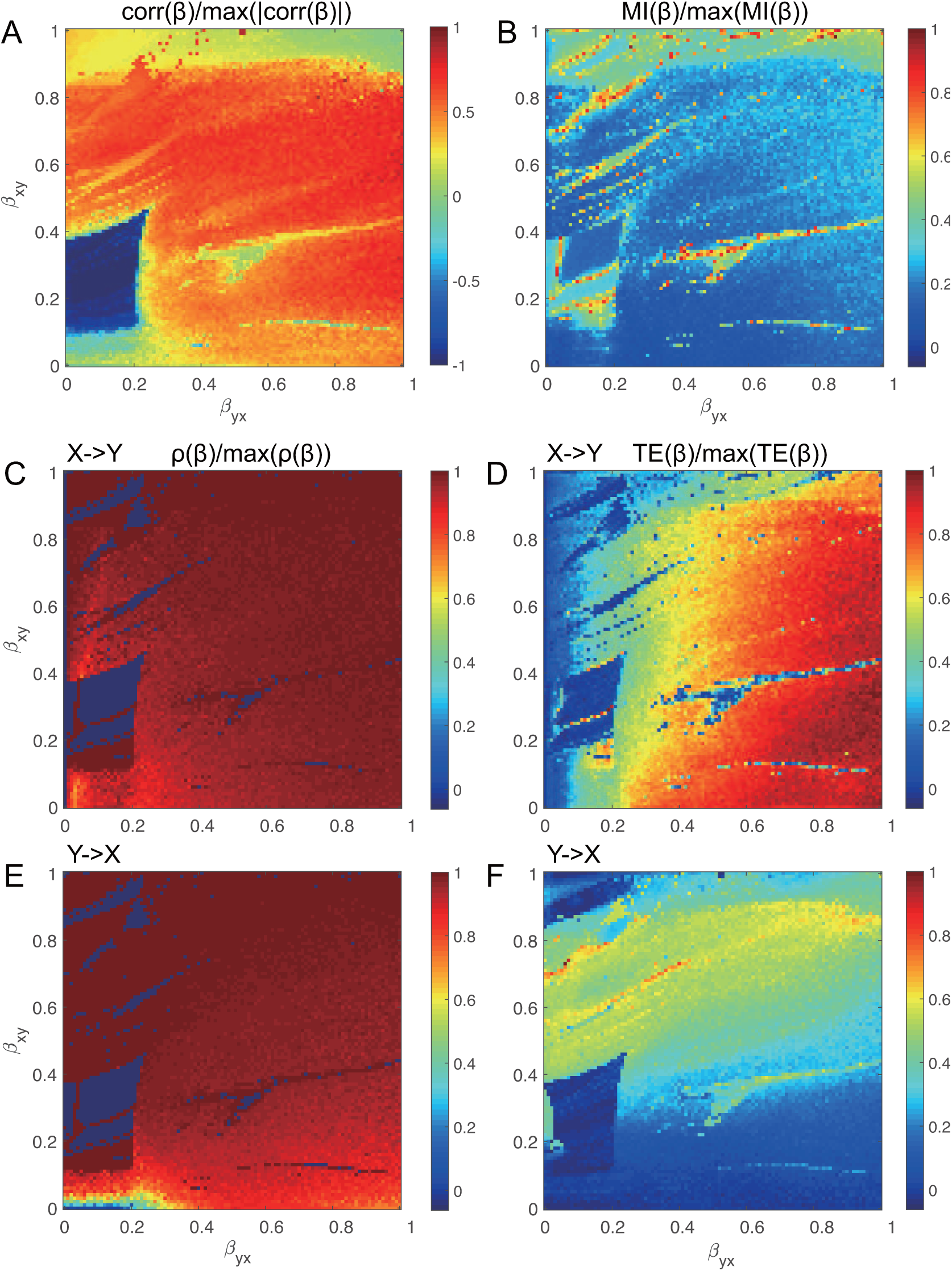
Phase-space maps of normalized coupling predictive causation via correlation, mutual information, CCM and OIF for varying true causal interactions. Both true causal interactions *β_xy_* and *β_yx_* are free varying within the range [0, 1], indicating a bivariate model S(*β_xy_*,*β_yx_*) where both species (or variables more generally) are interacting with each other with different strength. (A) normalized correlation coefficient; (B) normalized mutual information; (C) and (E) normalized CCM correlation coefficient (*ρ*) for interaction directions of *X → Y* and *Y → X*; (D) and (F) normalized transfer entropy (*T E*) from OIF model for interaction directions of *X → Y* and *Y → X*.

In a biological sense TE should be interpreted as the probability of likely uncooperative dynamics (leading to or driven by environmental or biological heterogeneity) while MI as the probability of cooperative dynamics (leading to or driven by homogeneity). Here we refer to cooperative and uncooperative interactions based on the similarity or dissimilarity in pair dynamics manifested by species abundance fluctuations. For instance divergence and asynchronicity (that define TE) in pair species dynamics manifest uncooperative interactions. The balance of cooperative and uncooperative interactions can result into net interactions at the ecosystem scale manifesting neutral patterns, or net interactions may lead to niche patterns biased toward strong environmental or biological factors^52^. Certainly, cooperation in a biological sense should be interpreted on a case by case basis. In a broader uncertainty propagation perspective ^49^, “cooperation” between variables means that variables contribute similarly to the uncertainty propagation, while “competition” means that one variable is predominant over the other in terms of magnitude of effects since TE is proportional to the magnitude rather than the frequency of effects. For the former case the total entropy of the system is higher than the latter case. Interestingly, correlation corr(*β*), *ρ* and TE show similar patterns in both organization and value range (but not in singular values of course), which sheds some important conclusions about the similarity and divergence of these methods as well as their capacity and limitations in characterizing non-linear systems.

When comparing the phase-space patterns from CCM and OIF (displaying *ρ* and *T E*) a more colorful and informative pattern is revealed by OIF. This means that TE gives a better gradient when tracking the increasing strength of causality for increasing values of *β_xy_* and *β_yx_*. When comparing the phase-space patterns for the two causal directions of “*X → Y*” and “*Y → X*”, phase-space maps from CCM are very similar, while those from TE present apparent differences in the strength of effects for the two opposite direction of interaction. Therefore, OIF is more sensitive to the direction of interaction compared to CCM when detecting directional causality.

These results imply that TE performs better to distinguish directional embedded physical interactions (that are dependent on direct interactions *β*-s, species growth rate *r_x_* and *r_y_*, and contingent values *X*(*t*) and *Y* (*t*) determining the total interaction as seen in the model of Eq. 5.1) in the species causal relationships. It should be emphasized how all linear and non-linear interaction indicators are inferring the total interaction and not only those exerted by *β*-s. In a broad uncertainty purview^49^ the importance of these three factors (*β*-s, *r*-s and *X*(*t*)/*Y* (*t*)) depends on their values and probability distributions that define the dynamics of the system; dynamics such as defined by the regions identified by patterns in Fig. 3 for the predator-prey system in Eq. 5.1. In principle, the higher the difference between these three interaction factors in the species considered, the higher the predictability and sensitivity of OIF. Fig. 4 highlights three different dynamics corresponding to the TE blue, green and red regions in Fig. 3.

**Figure 4.**
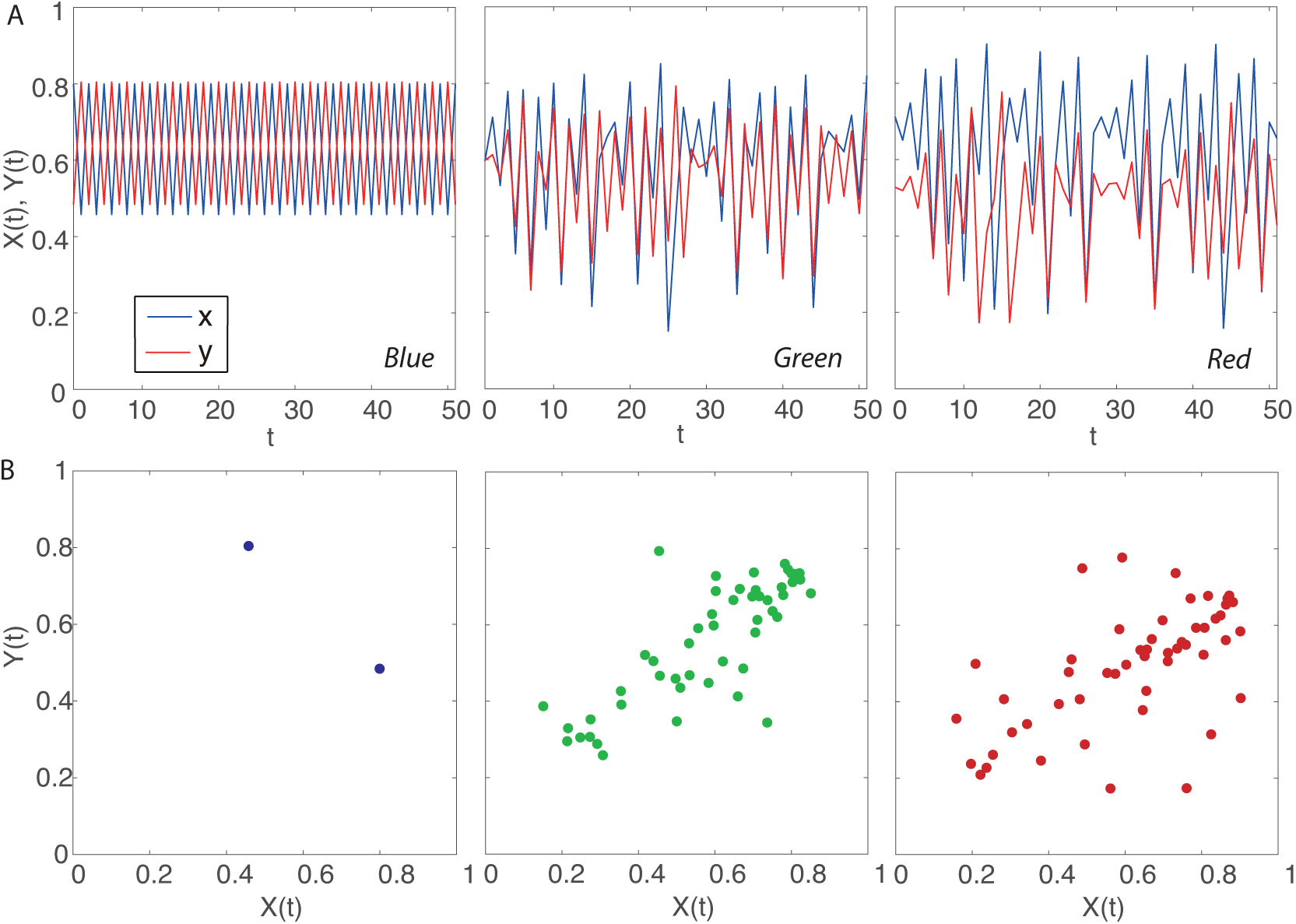
Dynamics of abundance and predictability for the bidirectional two species ecosystem model. (A) plots refer to the species abundance in time for the mathematical model in Eq. 2.1 for different predictability regimes associated to different interaction dynamics from low to high complexity ecosystem associated to low and high predictability. Blue, green and red refer to a range of predictable interactions as in Fig. 3: specifically, Blue is for (*β_y_ x*, *β_x_y*)=(0.18, 0.39) (small mutual interaction, and predominant effect of Y on X), Green is for (0.64, 0.57) (high mutual interactions, and slightly predominant effect of X on Y), and Red for (0.94, 0.34) (high mutual interactions, and predominant effect of X on Y). (B) phase-space plots showing the non-time delayed associations between X and Y corresponding to synchronous and homogeneous, mildly asynchronous and divergent, and asynchronous and divergent dynamics. The transition from synchronous/small interactions to asynchronous/high interaction lead to a transition from modular to nested ecosystem interactions when more than one species exist (Fig. 6).

In all dynamical states represented by Fig. 3, species are interacting with different magnitude and this defines distinct network topologies. Three prototypical dynamics are show in Fig. 4 with colors representative of *ρ* and TE in Fig. 3. The “blue” deterministic dynamics has very high synchronicity and no divergence considering variable fluctuation range (the gap is deterministic and related to the numerically imposed *u* = 1), as well as no linear correlation between non-lagged variables. In perfect synchrony one would have one point in the phasespace. Thus, absence of correlation does not imply complete decoupling of species but it can be a sign of small interactions. The “green” dynamics shows a relatively high synchronicity and medium divergence. In the phase-space of synchronous values of X and Y a correlation is observed with relatively small fluctuations because the divergence is small. Lastly, the “red” dynamics shows a relatively high asynchronicity and divergence. The stochasticity is higher than previous dynamics and the “mirage correlation” in the phase space has higher variance. Time-dependent mirage correlations in sign and magnitude mean that correlation (that may suggest common dynamics in a linear framework) does not imply similarity in dynamics for the two species. Non-linearity is higher from blue to red dynamics as well as predictability but lower absolute information entropy. Then, it is safe to say that linear dynamics (or small stochasticity) does not imply higher predictability.

### 2.3 Real-world Sardine-Anchovy-Temperature Ecosystem

CCM and proposed OIF model are also used for a real-world fishery ecosystem to infer potential causal interactions between Pacific sardines (*Sardinops sagax*) landings, Northern anchovies (*Engraulis mordax*) and Sea-Surface Temperature (SST) recorded at Scripps Pier and Newport Pier, California. Sardines and anchovies do not interact physically (or the interaction is low in number), while both of them are influenced by the external environmental SST that is the external forcing. To quantify the likely causal interactions between species and SST based on real data, we use CCM considering the length of time series for convergence of *ρ*, as well as OIF considering a set of time delays for acquiring stable values of inferred interactions TEs.

Results from CCM in Fig. 5A (plots from top to bottom) show that no significant interaction can be claimed between sardines and anchovies, as well as from sardines or anchovies in the SST manifold which expectedly indicates that neither sardines nor anchovies affect SST. This latter results, considering its biological plausibility should be taken as one validation criteria of predictive models, or complimentary as a test for anomaly detection of spurious interactions. The reverse effect of SST on sardines and anchovies can be quantitatively detected with the correlation coefficient *ρ* as well as TE. Although the calculated causations between SST and sardines or anchovies are moderate, CCM is able to provide a good performance in causality inference when the length of time series used is long enough due to convergence requirement.

**Figure 5.**
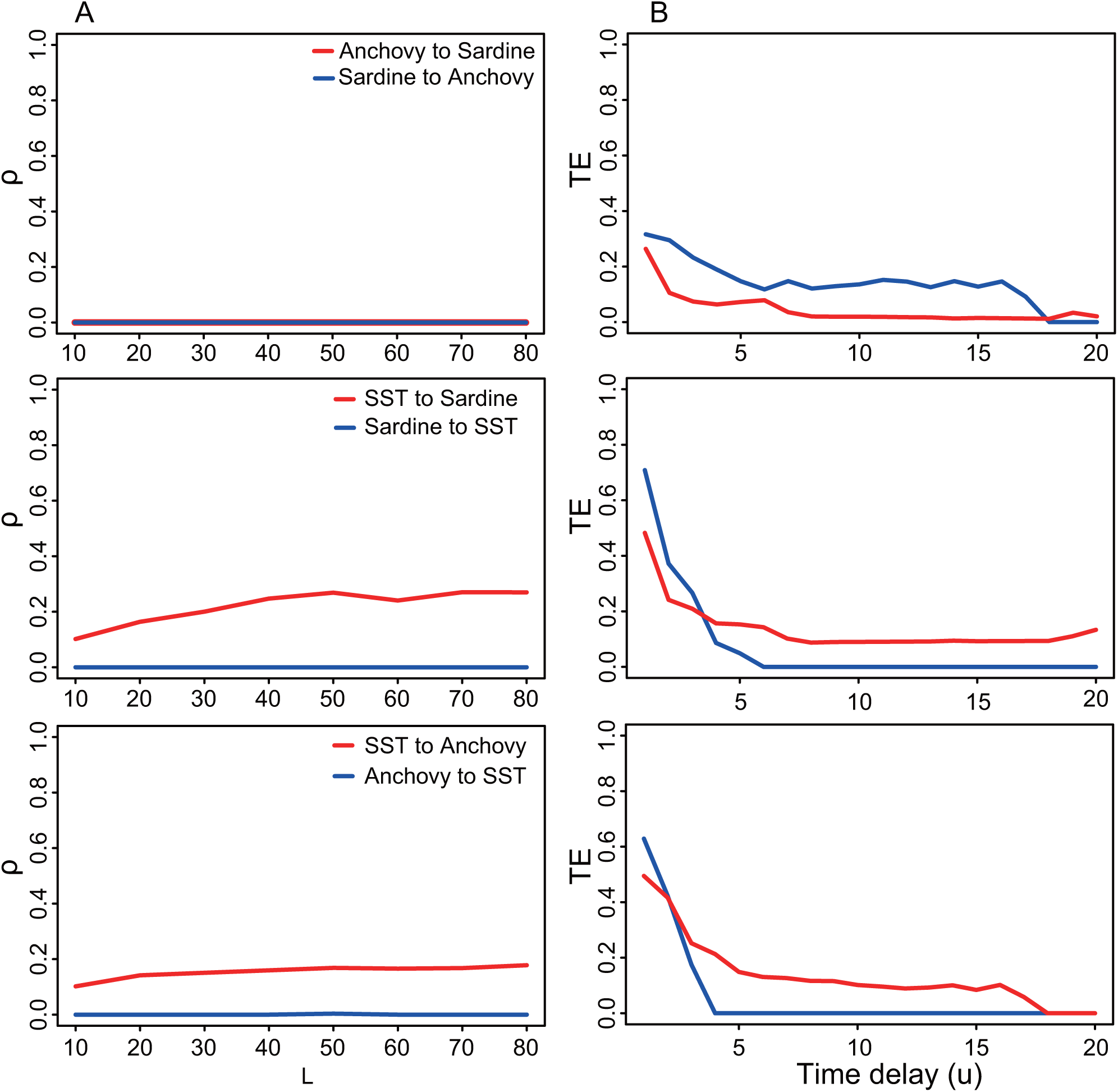
Inferred predictive causality for the sardine-anchovy-Sea Surface Temperature ecosystem. CCM correlation coefficient (*ρ*) and OIF predictor (TE) are shown in the left and middle plots for different pairs considered (sardine-anchovy, sardine and SST, anchovy and SST from top to bottom).

Figure 5B shows OIF’s results of inferred causal interactions between sardines, anchovies and SST dependent on the time delay *u*. For sardines and anchovies, OIF exposes bidirectional interactions that are actually biologically plausible, especially when both populations coexist in the same habitat, versus the results of CCM that infer *ρ* = 0. Ecologically speaking, even though fish populations do not directly influence sea temperature, we can find some clues about SST in fish populations influenced by SST. These clues can be interpreted as information of SST encoded in fish populations over abundance time records. So, observations of fish populations can be used to inversely predict the change of SST; this can be interpreted as “reverse predictability” (or “biological hindcasting”) in a similar way of when predicting historical climate change from ice cores. This information is captured by OIF, leading to nonzero values of TE from fish populations to SST. In this regard, we emphasize the distinction between direct and indirect (reverse) information flow, where direct information flow is most of the time larger and signifies causality (e.g. of SST for sardine and anchovies), and indirect (reverse) information flow that is typically smaller and signifies predictability (e.g. sardine and anchovies for ocean fluctuations). It is possible – especially for linear systems where an effect is observed immediately after a change – that information of SST encoded in fish populations is high if the interdependence, represented by the functional time delay *u*, of the environment-biota is small. However, for highly non-linear systems such as fishes and the ocean, changes in temperature may take a while before being encoded into fish population abundance ^53^. Thus, it is correct that the highest values of TE are for high *u*. Values of TE for small *u*-s are numerical artifacts related to systematic errors. Thus, for the effect of external SST on sardines and anchovies, OIF model gives unstable causal interactions with bias for lower time delays due to known dependencies of TE on *u* (such as cross-correlation for instance) that establishes the temporal lag on which the dependency between X and Y is evaluated. In a sense, plots in Figure 5B are like cross-variograms for the pairs of variables considered. TE becomes stable when the time delay is located in an appropriate range. It means that OIF requires an optimal time delay that makes results of the causality inference robust and that is related to optimal TEs (as highlighted in Li and Convertino (2019) ^39^ and Servadio and Convertino (2018) ^49^) that defines the most likely interdependency between variables for the *u* with the highest predictability. The fact that TE of sardine and anchovies to SST is high for same small ranges of *u* may be also a byproduct of data sampling, i.e., fish and SST sampling locations are different (fish abundance is actually about fish landings) and that can introduce spurious correlations/causation. Overall, these findings suggest that the OIF model provides more plausible results, but it requires careful selection of optimal time delays.

Figure S1 shows the relationships between normalized *ρ* and TE estimated for all selected values of *L* and *u* of pairs in Fig. 5 (sardine-anchovy, sardine and SST, anchovy and SST). These plots show opposite results than the proportionality between *ρ* and TE in Fig. 3 because non-optimal values are used, that is non-convergent *ρ*-s and suboptimal TE during the interaction inference procedure (Fig. S1). TE for too small *u*-s determines overestimation of interactions due to the implicit assumptions that variables have an immediate effect on each other and that is not always the case as highlighted by the vast time-lagged determined non-linear regions in Fig. 3. If “transitory” values of *ρ* for small *L* are disregarded, as well as TEs for small *u*-s, the relationship between *ρ* and TE shows a correct linear proportionality.

### 2.4 Real-world Multispecies Ecosystem

Interactions between fish species living in the Maizuru bay are intimately related to external environmental factors of the ecosystem where they live, the number of species living in this region considering also the unreported ones) and biological species interactions, which leads to a complex dynamical nonlinear system. In Fig. 6 the network of observed fish species (Table S1) is reported where only the interactions considered in Ushio et al. (2018) ^37^ for the CCM are reported. This is because the goal is to compare the CCM inferred network to the TE-based one based on abundance. Fig. 7 shows the temporal fluctuations of abundance and the functional interaction matrices of *ρ* and TE. In this paper we study and compare average ecosystem networks for the whole time period considered but dynamical networks can also be extracted via time-fluctuating *ρ* and TE as shown in Fig. S3. These dynamical networks can be useful for studying how diversity is changing over time and ecosystem stability (Figs. S4, S6-S7) as well as understanding the relationship between *ρ* and TE (Fig. S5). In the network of Fig. 6 the color and width of links is proportional to the magnitude of TE (Table S2); for the former a red-blue scale is adopted where the red/blue is for the highest/lowest TEs. The diameter is proportional to the Shannon entropy of the species abundance pdf (Table S3). The color of nodes is proportional to the structural node degree, i.e. how many species are interconnected to others after. Therefore, the network in Fig. 6 is focusing on uncooperative species whose divergence and/or asynchronicity (that is a predominant factor in determining TE over divergence) is large. Yet, the connected species are rarely but strongly interacting in magnitude rather than frequently and weakly (i.e., cooperative or similar dynamics). Additionally, the species with the smallest variance in abundance are characterized by the smallest Shannon entropy (smallest nodes) and more power-law distribution although the latter is not a stringent requirement since both pdf shape and abundance range (in particular maximum abundance) play a role in the magnitude of entropy. Average entropy such as average abundance are quantities with limited utility in understanding the dynamics of an ecosystem as well as ecological function. Nonetheless, species with high average abundance (e.g. species 5) have a very regular seasonal oscillations and the largest number of interactions with divergent species. A result that is expected considering the size of the population and the ability of the species to follow regular environmental fluctuations.

**Figure 6.**
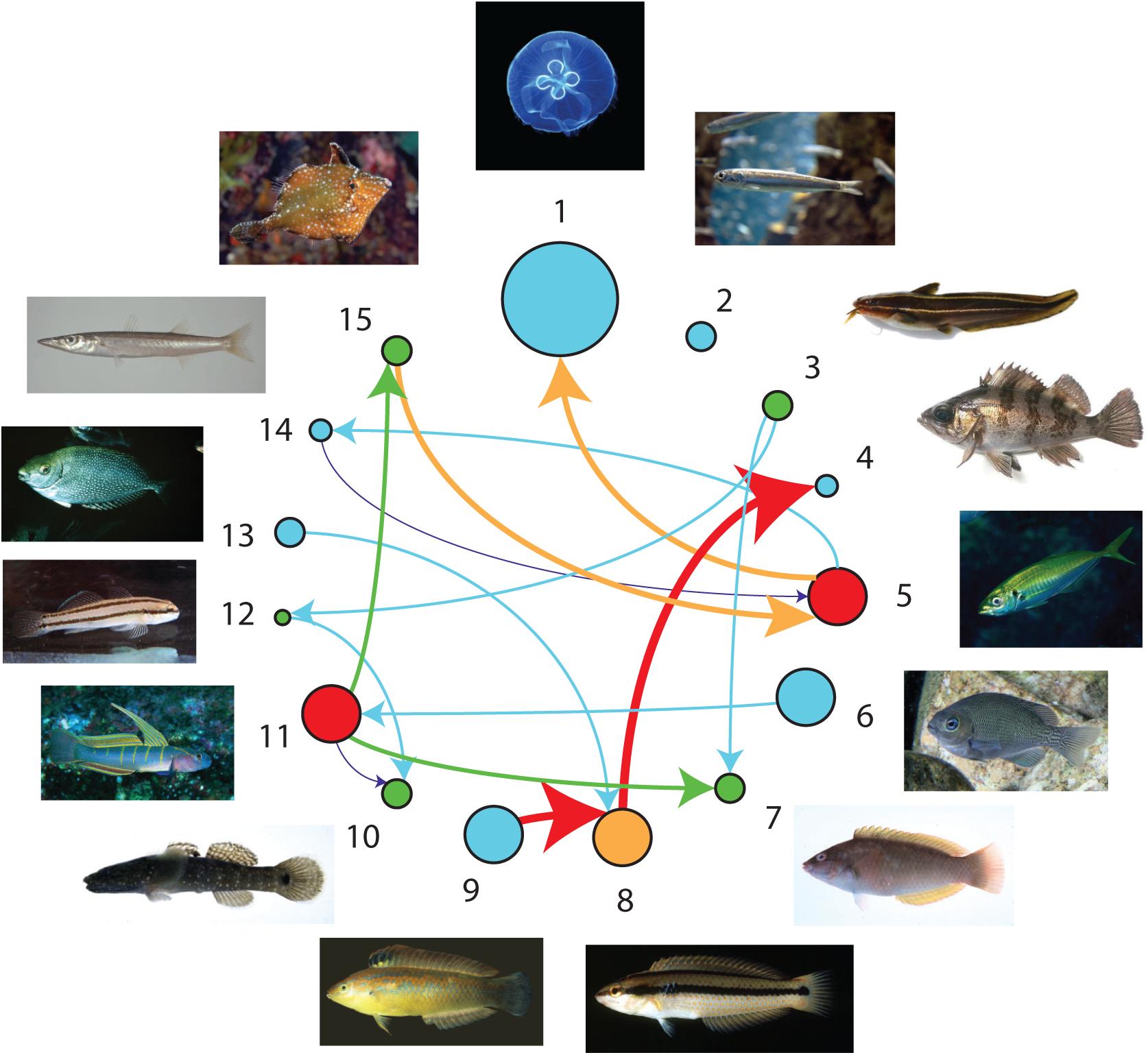
Part of the estimated species interaction network for the Maizuru Bay ecosystem. Species properties are reported in Table S1. The color and width of links is proportional to the magnitude of TE (Table S2); for the former a red-blue scale is adopted where the red/blue is for the highest/lowest TEs. The diameter is proportional to the Shannon entropy of the species abundance (Table S3) that is directly proportional to the degree of uniformity of the abundance pdf and the diversity of abundance values (e.g., the higher the zero abundance instances the lower the entropy). The color of nodes is proportional to the structural node degree, i.e. how many species are interconnected to others after considering only the CCM derived largest interactions (see Ushio et al. (2018) ^37^ and Fig. 7). Other interactions exist between species as reported in Fig. 7. TE is on average proportional to *ρ* (Fig. S4 and S5).

**Figure 7.**
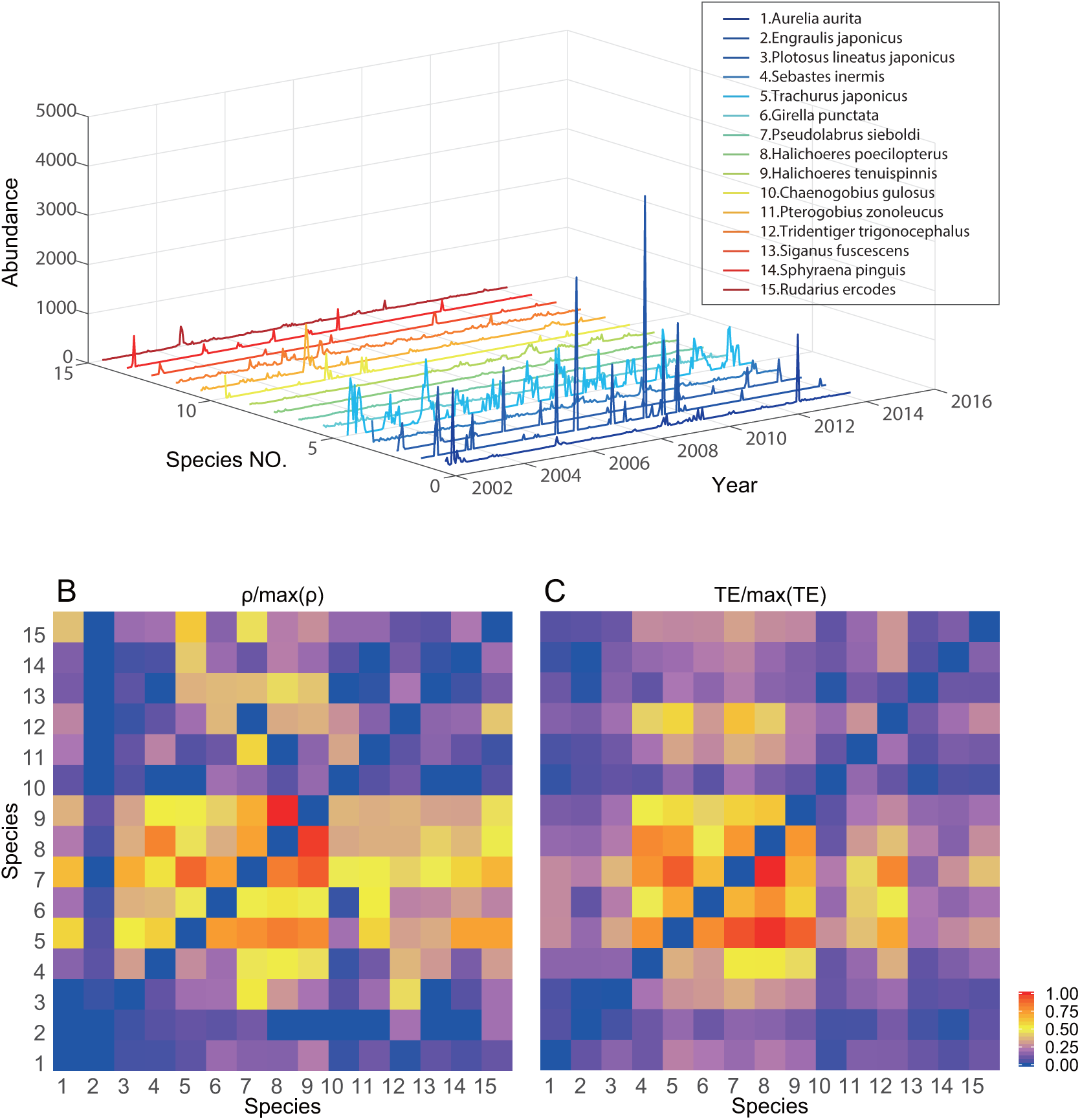
Normalized species interactions matrices inferred by CCM and OIF models for Maizuru Bay ecosystem. In the census of the aquatic community, 15 fish species were counted in total. Interaction inferential models use time lagged abundance magnitude (CCM) or pdfs of abundance (OIF) shown in A. (B) normalized CCM correlation coefficients (*ρ*) between all possible pairs of species. (C) normalized transfer entropies (TEs) between all pairs of species from the OIF model. Both CCM and OIF predict that the most interacting species (in terms of magnitude rather than frequency) are 7, 8 and 9 on average. Thus, interaction matrices are more proportional to the asynchronicity than the divergence of species in terms of abundance pdf, although abundance value range defines the uncertainty (and diversity) for each species that ultimately affects entropy and interactions (e.g., if one species have many zero abundance instances or many equivalent values, such as species 2, TEs of that species are expected to be low due to lower uncertainty despite the asynchrony and divergence).

Figure S2 shows that the strongest linear correlation is for the most divergent and asynchronous species (from species 4 to 9) for which both *ρ* and TE are the highest (Fig. 7 B and C). This confirms the results of Fig. 3 and the fact that competition (or dynamical diversity more generally) increases predictability. This also highlights the fact that linear correlation among state variables does not imply synchronicity or dynamic similarity as commonly assumed. The interaction matrices in Fig. 7 B and C confirm that TE has the ability to infer a larger gradient of interactions than *ρ* and the total entropy of the TE matrix is lower than *ρ*. Pairwise the inferred interaction values by CCM and OIF are different but *ρ* and TE patterns appear clearly similar and yet proportional to each other.

CCM and OIF models are applied to calculate the potential interactions between all pairs of species. Fig. 7B and C show interaction matrices describing the normalized *ρ* from CCM and TE from OIF model of all pairwise species, respectively. The greater the strength of likely interaction, the warmer the color. These results demonstrate that CCM and OIF model present similar patterns for the interaction matrices in terms of interaction distribution, gradient and magnitude in order of similarity. This indicates that both CCM and OIF are able to infer the potentially causal relationships between species. Compared to the CCM interaction heatmap the OIF heatmap presents larger gradients of inferred interactions that highlight the divergence and asynchrony in fish populations of species 4-9 from other species. This difference can be observed in Figure S2 that shows the strongest linear correlations for the most divergent and asynchronous species (4-9). It is worth noting that species 4-8 are all native species (See Table S1). Therefore, despite the patterns of interactions of CCM and TE are similar, CCM allows one a better identification of clusters of species with similar or distinct interaction ranges. Additionally TE estimates some weak observed interactions such as of species 2 (*E. japonicus*) with others, while CCM essentially consider null interactions for these species.

Precisely, the most interacting species (4-9) are the most divergent and asynchronous species (with respect to the whole community) as well as diverse in terms of values of abundance; these species form the “collective core” that is likely determining the stability of the ecosystem. Interestingly the number of these species is relatively small and it confirms results of other studies (see ^54;55;56;57;39^) showing that the number of species with weak interactions is much larger. Theory suggests that this pattern promotes stability as weak interactors dampen the destabilizing potential of strong interactors^54^. Mediated cooperation (e.g. by many “weakly” interacting competitors) as shown by Tu et al (2019) ^52^ promotes biodiversity and diversity increases stability. When considering abundance values (at same time steps) of collective core species (Fig. S2) these species are linearly related and this increases their mutual predictability by either using linear or non-linear models based on correlation coefficient and TE. This proves that non-linearity increases predictability.

The choice of the optimal *u* that maximizes MI leads to the optimal TE model and resultant interaction network. The observed *u* over time is really small (Fig. S9) and this signifies how likely the ecosystem has small memory and responds quickly to rapid changes, or the information of change is carried over time by ecosystem’s interactions which leads to accurate short-term forecasting. In other words, temperature-induced changes may take long time but the information of change is replicated at short time periods. The chosen time delay *u* = 1 corresponds to the species sampling of two weeks. Note that values of *u* are also dependent on the data resolution and they are strongly related to fluctuations rather than absolute *α*-diversity value. Thus, while biodiversity may fluctuate rapidly in time, value of *α*-diversity for seasons or longer time periods can be more stable and manifesting higher memory (representative of *u* for the whole ecosystem) than the one between species pairs (related to pair’s *u*). Short-term catastrophic dynamics (for instance related to dramatic habitat change, sudden invasions, extinctions or rapid adaptations) may lead to irreversible shifts in interactions (strength and sign); this, in turn can affect biodiversity patterns that are completely uninformed by past dynamics. Thus, there is certainly a limit to predictability and to the validity of time delays which can change very rapidly. However, we insist in emphasizing that models are *predictive tools* and predictions is not necessarily causality reflecting the many and highly complex underlying processes. Yet, interpretation of results must be done with care.

We also study temporally dynamical networks for the fish ecosystem community (see Section 2.3). CCM and OIF model are applied to quantify the causality between all possible pairs of species at each time period by calculating *ρ* and TE, respectively. Estimated effective *α*-diversity (Eq. 2.8) from CCM- and TE-based inferred networks at each time point can be obtained and then compared to the taxonomic (or “real”) *α*-diversity. Results are shown in Fig. 8 and Fig. S6. In the whole time period, the estimated *α*-diversity from CCM is constant, whereas the global trend of the estimated *α*-diversity from OIF model slightly decreases over time that is consistent with the global trend of real *α*-diversity. CCM always predicts a non-zero interaction for all species (including negative values) whereas OIF predicts zero interactions for some species that are then not making part of the estimated effective *α*-diversity.

**Figure 8.**
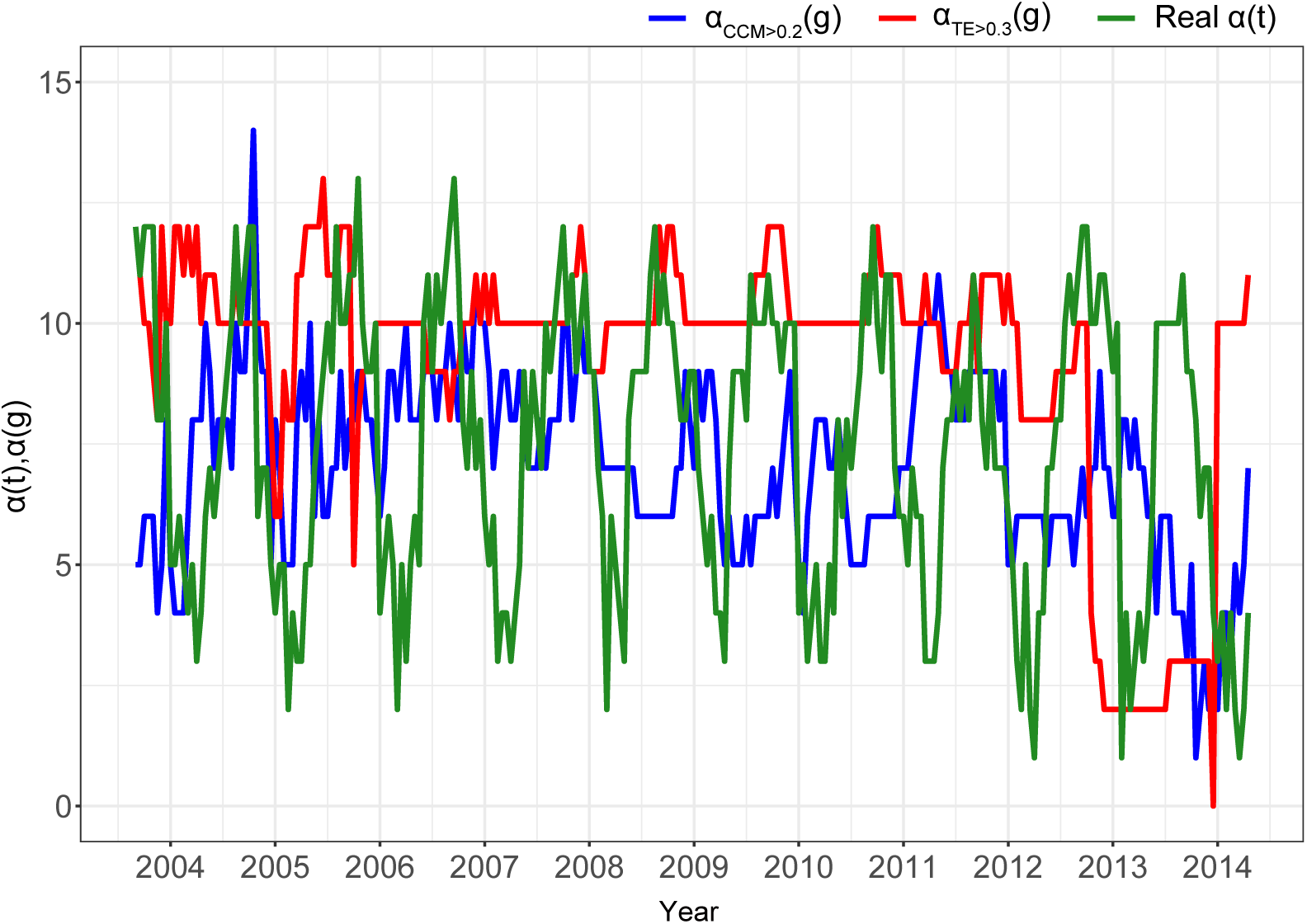
Predicted α-diversity via optimal interaction threshold for CCM’s ρ and OIF’s TE versus taxonomic diversity. Effective *α* diversity from CCM and OIF are shown (blue and red) for an optimal threshold of *ρ* and TE (i.e., 0.2 and 0.3) that maximizes the correlation coefficient and Mutual Information (MI) between *α_CCM_* or *α_TE_* and the taxonomic *α*, respectively. The maximization of the correlation coefficient and MI guarantees that the estimated effective *α* are the closest to the taxonomic *α*. *g* is the resolution of the network inference determined by the minimum number of points required to construct pdfs and infer TE robustly (see Supplementary Information section S1.3).

Figure 8 shows the effective *α* diversity from CCM and OIF for an optimal threshold of *ρ* and TE (i.e., 0.2 and 0.3) that maximizes the correlation coefficient and Mutual Information (MI) between *α_CCM_* or *α_TE_* and the taxonomic *α*, respectively. The maximization of the correlation coefficient and MI guarantees that the estimated effective *α* are the closest to the taxonomic *α*. Figure S6 shows effective *α* for unthresholded interactions and other thresholds. Note that the threshold on TE does not coincide with the value of TE that maximizes the total network entropy (Fig. 8) and then some of the reported species may not be part of the ecosystem strongly. Thus, this threshold method is also useful to identify species that are truly forming local diversity vs. transient species. Considering the pattern of fluctuations of effective *α*-diversity from CCM they are poorly unrelated to the real *α*-diversity, while those from OIF are much more synchronous with seasonal fluctuations of real *α*-diversity. However, *α_TE_* is a bit higher than the average taxonomic *α*. Both CCM and TE predicts a decrease in *α* in time that corresponds to an increase in SST. As shown in Fig S6, OIF is attributing higher sensitivity to SST for small interaction species because *α* fluctuations show seasonality that happens when species follow environmental dynamics closely. Vice versa, CCM is predicting a broader sensitivity for all positively interacting species. These results reveal that OIF gives an effective tool to measure meaningful interdependence relationships between species for constructing temporally dynamical networks where the number of nodes over time (estimated *α*(*t*)) can reflect closely the taxonomic *α*-diversity. This allow us to find more reliably how changes of environmental factors (e.g. SST) affect biodiversity in ecosystems. The establishment of thresholds on interactions is also useful for exploring ranges of interdependencies and associated effective *α*-diversity with respect to the average taxonomic diversity. *β* effective diversity is another very important macroecological indicator informing about ecosystem changes; for instance in Li and Convertino (2019) ^39^ *β*-diversity identified distinct ecosystem health states. However, *α* and *β* (effective diversity) variability are highly linked to each other and yet looking into one or another would provide equivalent results. The difference between taxonomic and effective *β*-diversity may provide some information about invasive or rare species have weak influence on the ecosystem since they are characterized by low TEs. Supplementary Information contains further elaborations on results.

## 3 Discussion

In the paper the proposed Optimal Information Flow (OIF) model was validated by considering the problem of causality inference of species interactions for ecosystems with different level of complexity and systemic uncertainty: a deterministic mathematical model of predator-prey dynamics, the real Sardine-Anchovy-Temperature triplet ecosystem in the Pacific, and a real multispecies fish ecosystem in Japan. These three case studies are epitomic example of deterministic, low and high complexity dynamics. The mathematical model can be generalized as a model for interaction dynamics between individual or communities of the same species or between two generic variables X and Y. The quantification of interactions was compared to the well-documented CCM model.

Early method of correlation were proved to be neither necessary nor sufficient to estimate the causal relationship between time series variables (mostly due to the fact that any association does not prove causality because models are not surrogate of reality and scale-dependent data are just a sample of ecosystem dynamics), even though it remains a common and heuristic notion^35 29^. Despite these views we prove the power and limitations of correlation methods with respect to non-linear methods (such as OIF and CCM), dynamics of the complex ecosystem considered, and target patterns to predict that define whether correlation is able to measure interactions. As for the latter, for instance we show how even highly non-linear systems show linearity when non-linear “causal” variables are considered at the same time step (as a virtue of non-linear asynchronicity). Therefore, the scale of analysis considering also the space-time domain with the explicit consideration of lag effects, determines the dynamics that is visible and the model that can be used for predictions.

Granger causality is the primary framework that uses predictability especially for identifying causation, however it is problematic in highly nonlinear systems even with some deterministic states or components. Sugihara et al. (2012) ^29^ managed to deal with the problem and introduced the CCM model. CCM was well documented and successfully applied to bio-inspired mathematical models, as well as real-world ecosystems ^29^. Despite interactions among species or variables, or interdependencies more generally defined, are rarely completely zero and related to patterns of different processes to capture, Sugihara et al. (2012) ^29^ maintains the view of a deterministic single-value causality. In our opinion calculating causality in an absolute sense between variables is always not only very hard, but also meaningless because the resulting values are dependent on data and models used as well as the predicted patterns for which interactions are calculated for. The very first question should be causality about what? After that the evaluation of the dynamics of the ecosystem coupled to the target patterns to map should drive model selection. Causality is actually predictability of patterns of interest and predictability can be close to true causality for systems with low complexity and noise. The basic principles to interpret predictability are uncertainty reduction and accuracy that can be quantified as the probability of an event to occur given another one (as predictands and predictors, respectively).

From the perspective of information theory that has attracted attention in complex networks research, entropy is the information-theoretic description for uncertainty or more precisely lack of organization rather than absolute uncertainty. Uncertainty is in fact also information about the diversity of values of a complex system (see e.g. Jost (2006) ^50^ that demonstrated how entropies are reasonable indices of diversity) and the distribution of these values determines entropy. The fundamental work of studying complex networks is to untangle complex interdependencies comprising a large number of potential causations between all pairwise nodes (variables), that allows one to predict the collective behavior of complex systems. The intuitive and heuristic notion for this problem in information theory is transfer entropy that measures the uncertainty reduction (or information flow) between nodes (variable). From this conceptual perspective we name the OIF model for inferring potential causality seen as sets of uncertainty reduction networked fluxes. Multiple transfer entropies for one single variable as a function of all others determine non-linearity that cannot be overlooked even when variable interaction is deterministic. Considering entropy as diversity also implies that OIF provides reflections of temporal changes in diversity (e.g. biodiversity) determined by changes in information fluxes.

The bio-inspired mathematical model generates a clean inter-species interaction ecosystem without any noise, that allow us to estimate “true” causality between synthetic species X and Y. The so-called “true” causality means the causation embedded numerically in the parameters in the dynamical equations 2.1 (*β_xy_* and *β_yx_*). When *β_xy_* is fixed as zero only the unidirectional causality (*X → Y*) exists between species X and Y. Then, any estimator of predictive causality closer to the physical causality *β* defines the accuracy of the model. Results from Fig. 2 shows how OIF model outperforms CCM.

Depending on the values of the parameters the model may capture some biological dynamics such as amensalism and commensalism (when *β_xy_* or *β_yx_* are zero), or predation, competition and mutualism (when both *β_xy_* and *β_yx_* are different than zero). Biologists define amensalism (i.e. a strong asymmetrical competition) is a type of biological relationship between species in which one species (e.g. X) has a potential negative effect on another (Y), but the second species Y has no detectable effect on the first species X. Biologically speaking, commensalism is another type of biological relationship in which one species (Y) gets benefits while the other one (X) is neither helped or harmed.

In a broad complex dynamic perspective it is easier to talk about “cooperation” or “competition” between variables meaning that variables contribute similarly to the uncertainty propagation of the whole ecosystem, or that one variable is predominant over the other in terms of magnitude vs. frequency of effects. Similarly to brain function^58^, cooperation is typically driven by excitatory interactions leading to synchronization in biomass fluctuations, where species are characterized by power-law distributions; vice versa competition is driven by inhibitory interactions leading to asynchronization, where species are characterized by exponential or non-fat tail distributions of abundance. The generic dynamical characterization allows to avoid pitfalls of the categorical classification of interactions in biology that suffers from the lack of knowledge about true and meaningful values of interactions that distinguish one biological dynamics from another; this is also considering change in biological interactions over time driven by environmental changes and/or evolution. Depending on the ecosystem state, interactions might range between “positive” and “negative” with no clear cuts between them: e.g. ^39^ showed that network topology and interactions among the same microbial species change for different ecosystem states. We also caution to use numerical estimates of interactions to replace empirical biological knowledge because data-inferred interactions are always much more complex than experimental values (e.g. about highly controlled lab tests) and interactions are certainly highly affected by the environmental context, measurement technology, and biomarker considered (e.g. abundance or others). The similar pattern of inferred interactions of the predator-prey system shows that all methods (correlation, MI, CCM and OIF) can work for inferring causality between two variables with different level of granularity. However, considering our definition of predictive causality, as non-linear predictability of diverse events from independent predictors, OIF outperforms all other models due to the explicit consideration of asynchronicity, divergence and diversity of events that define model sensitivity, uncertainty and complexity. All these considerations further emphasize the need to distinguish biological interactions and model-driven inferred interdependencies (or predictive interactions) as also emphasized by other authors^51^.

To analyze OIF performance for “low complexity” ecosystems we considered the ambiguous dynamics of sardines and anchovies in oceans. On multidecadal time scales, sardines and anchovies present alternating dominance across global fisheries. Although in in appearance a ecological competition seems to exist between these two species (due to the inversely proportional and synchronized abundance changes), the simultaneous fluctuations of sardine and anchovy stocks suggest that they are also influenced by the ocean temperature.

Incompatible hypotheses have been advanced to try to give explanations for this pattern of alternating dominance, unfortunately leaving aside many other species that clearly exist in the ocean and interacting with sardines and anchovies. Some supposed that these two species act in direct and clear competition^59^, while others argued that this pattern is just a result of different or opposite fish dynamics in response to common global environmental forces ^60^. Results in Baumgartner (1992) ^61^ revealed that in longer time series not only the negative cross-correlation observed in the 20th century disappears, but the correlation with global environmental forces also has been ambiguous. This lack of correlation is however only related to the fact that species are synchronous and environment*→*species effects are characterized by relatively small lags. Yet, lack of evident correlation exist but that does exclude causation. Jacobson and MacCall^62^ applied two models to this issue and proposed a relationship that SST influences the behavior and population of sardines and anchovies; however, this relationship vanished when applying the analysis to stock assessments from 1992 to 2009. Although all these possible explanations from different points of view are competing, or even unstable, such results can illustrate that causal interactions among sardines, anchovies and SST present features of nonlinear dynamics. Nonetheless, and more importantly, the conclusion is that both species are weakly interacting and majorly affected by the environment. All interdependencies exist and they just change in terms of normalized magnitude without neglecting the fact that intrinsic interspecies interaction is also modulated by the environment. As shown by the predator-prey mathematical model (Fig. 3) and real data (Fig. 5 and 7), synchronized species are certainly affected by a third variable (e.g. the environment and other species) that is forcing both in fluctuating at the same time.

Heuristically, it is also very unlikely that two species (or variables more generally) are perfectly synchronized unless they are identical. What Fig. 4 shows is somewhat very affine to the Heisenberg’s uncertainty principle that marks a clear break from the classical deterministic view of the universe. We cannot know the present state of the world in full detail (such as for the “red” dynamics), let alone predict the future with absolute precision. Determinism, driven by synchrony, allows us to know the current state of the system if that is unaltered but not to predict future states. Vice versa, uncertainty-driven asynchronicity and divergence allow us to predict likely future more than actual present and that appears to be in contradiction to deterministic views but not to realistic probabilistic (or relativistic) view of system dynamics. For this sardines-anchovies “problem” unfortunately the whole complexity of ecosystems has never been considered despite other species may have a dominant effect on their abundance. This underlines the importance of space-time and biological scale (where biological scale is define by the number of interdependent species at the same or different trophic levels) in framing the problem and bounding conclusions to model results: any “causation” is in reality an interdependence between species constrained to the chosen scales, data resolution, as well as model analytics and biomarkers used for inferring interdependencies. For instance, lack of large environmental disturbances (or coarse resolution sampling) affecting rapidly small organism change may reduce the information flow, and yet affecting the inferred interaction network.

In addition to testing OIF on the simplified sardine-anchovy ecosystem, we apply OIF and CCM to a multispecies ecosystem in which 14 dominant fish and 1 jellyfish species were monitored in an abundance census in the Maizuru Bay, Japan. In this ecosystem, all species can be interconnected, leading to an intricate causality system that is extremely hard to estimate considering intrinsic biological species interactions, interactions related to environmental influence, and biological interaction mediated by the environment. “True causality” assessment is also extremely hard because there is no knowledge of which ecological or biological marker can capture all these interaction types. However, when causality is shifted to predictability of patterns of interest, the issue of inferring causality becomes practical and meaningful. Predictive causality between species X and Y, for instance, depends on whether X can assist in predicting the future of Y beyond the extent to which Y itself predicts its own future, and complementarily whether the model can predict the collective behavior of the system which can be reflected by macroecological indicators dependent on all predictive causality. In this case study, both CCM and OIF models are effective for causality detection from different points of view and they majorly differ considering interaction gradient and computational complexity. As for the latter OIF and CCM have 3 and 20 parameters to populate and the speed of inference assessment for the Maizuru 15 species ecosystem is 2 and 15 minutes, respectively. An already published work used the same time-series data to study how to infer the network and forecast the system stability for the fish community using CCM^37^.

Ushio et al. (2018) ^37^ used a “S-map” model (i.e., sequential locally weighted global linear map model) to track dynamical interactions over causally related species over time where causality between species is estimated a-priori via CCM. S-map does so by predicting future values of species abundance from the reconstructed multivariate state-space vector (where species are interdependent by inference). Before CCM a “phase-lock twin surrogate” method removes seasonality from abundance data; this method generates time series that preserve the shape of a species attractor but exhibit no causal relationship with a target (seasonality) time series. Other predictive models other than S-map have been proposed such as the multiview embedding model ^63^ (in a “Empirical Dynamic Modeling” suite of models ^64^) that is combining multiple species embeddings (with different time delays) and that is superior in forecasting skill than multivariate (e.g. S-map) and univariate embedding (used by CCM for reconstructing the attractor of each species individually and later on the cross-mapping skill *ρ* of each other is evaluated).

Here we believe that any environmental forcing is important to be captured and affect non-linearly and in an unpredictable way the interactions among species; pure biological interactions are utopianly impossible to measure (leading to the “curse of environment separability” from biota) and they are always context dependent (i.e. the geographical area considered although universality in biological dynamics may be expected). While it may be true that synchronization driven by seasonality can lead to misidentification of “biological” or “true” causality (false negative without the consideration of time lags, or false positive as in Ushio et al. (2018) ^37^ if lags are considered in the phase-space), we believe that the environment is precisely the common identifiable cause of synchronization of species (or of other asynchronous effects impacting species differently) in a predictive causality purview. Additionally our is a more realistic analysis of ecosystems where the environment is central in shaping interconnected populations, and then community patterns, via complex non-linear function vs. simple assumed sinusoidal seasonality (homogeneous for all species). Lastly, in our opinion another pitfall of the “phase-lock twin surrogate” model of Ushio et al. (2018) ^37^ is the fact that seasonality importance is weighted for each species in isolation whereas seasonality is also affecting interactions of species pairs in ecosystems. Interactions that arguably should be inferred from data as they are since data contain hardly separable non-linear effects of environment and other species dependencies in addition to single species adaptation and evolution.

OIF, through the inference of a better gradient of systemic interaction “causality”, predicts how biodiversity changes over time with average value, fluctuations and trend that is closer to the taxonomic *α*-diversity. This is for effective *α* diversity with the optimal threshold on interactions maximizing the similarity with observed *α* (via maximization of Mutual Information). The concept of effective *α* is very useful because it allows to see which set of interactions is determining levels of *α* diversity that is potentially more or less sensitive to environmental forcing. For example, S6 shows that high interaction species form a small portion of community diversity that is increasing over time vs. the systemic decrease in diversity (observed in Fig, 8). More importantly, the increasing fluctuations of estimated *α* from OIF show the potential way in which climate and/or other anthropogenic changes negatively affects biodiversity in the region considered in relation to intensified interspecies interactions as suggested also in other studies^65;66;67^. These results are certainly beneficial for fishery resources management and habitat protection aiming to preservation of the fish community with ecological, economic and social outcomes. Thus, models like OIF should be evaluated in this bigger perspective of ecosystem utility or ecological engineering with multiple utilities rather than just seeing these models for the hard inference of pure “biological” interaction causality. Supplementary Information contains further discussions about CCM, TE causality inference, predictability, ecosystem organization and stability.

## 4 Conclusions

Causality detection is a fundamental step in the inference of complex networks with the aim of understanding processes of observed complex systems. This is incredibly important for poorly observable large scale ecosystems whose structural and functional networks are their backbone. However, quantifying the “truly causal” interactions in complex systems is illusory and perhaps impossible to achieve due to data and model limitations (e.g. sampling over space and time), partial ignorance about underlying processes, the strong unmeasurable influence of environmental dynamics, and more importantly their relativity dependent on the scale of analysis and the patterns for which interactions are relevant for. Nonetheless, when causality is shifted to predictability, this issue becomes practical and useful because it links causal *predictable* interactions to some patterns to predict. Patterns that are defining the socio-ecological outcomes of interest for which interactions are signatures of the underlying processes. In this paper we propose the Optimal Information Flow (OIF) model and assess its validity and performance in causality inference by comparing OIF to well-documented CCM and correlation model. This is done for a deterministic predator-prey mathematical model, a data-driven sardine-anchovy species dynamics, and an observed multiple fish species ecosystem. We show that OIF, like CCM, is able to effectively identify asymmetric causal interactions between any pair of species. Moreover, OIF performs better than CCM because it provides: (i) a larger gradient of interaction values, yet defining interactions at higher resolution with better definition of asymmetrical interdependencies; (ii) smaller fluctuations around the estimated interaction values for any time delay *u*, yet a less uncertain inference; (iii) the estimated memory of one and pairs of species in terms of time delay (without considering future modifications of CCM^68^); (iv) independence on the length of historical data and no requirement for convergence, as well as lower computational complexity (leading to lower sensitivity and uncertainty in state estimates); and, (v) more accurate predictions of temporal changes in macroecological indicators of ecosystems such as for the effective *α*-diversity after optimal MI-based threshold selection. However, OIF requires the identification of the optimal *u* value as shown in^39^ but this is easily automated by exploring the delay that maximizes MI. Even though a time delay can be defined for any pair of species, we show the the average time delay, derived from analyzing all species pairs, can be a global optimum providing accurate macroecological predictions for the ecosystem considered. Thus, the assumption-free information-theoretic OIF is a strong candidate model for the inference of predictable causality in complex ecosystems. A model that is itself an ecosystem mimicking the information flow constituting the backbone of real ecosystems of any nature, from environmental to socio-technological systems. The complexity of real world systems might be higher than the ones studied in this paper, considering the velocity of transitions in rapidly changing systems. Nonetheless, we believe that the dynamics encompassed in our study reflects the fundamental stochastic processes observable in the real world, particularly at stationarity but changes in network topology can be mapped by inferring dynamical networks over time. In a broader uncertainty propagation perspective interactions should be considered as “cooperation” and “competition” between species (or variables more generally) meaning that they contribute in a similar or opposite way to the uncertainty (or information) propagation. Competition means that one variable is predominant (or very diverse) over the other in terms of magnitude of interactions since TE is proportional to the magnitude rather than the frequency of interactions. Interactions that are specifically proportional to the divergence and asynchrony of variables/species which leads to higher predictability. In conclusion, our model can find useful applications in research and applied work for ecosystems at multiple biological scales. A myriad of other models have been proposed in literature, and these can be used simultaneously in real-life applications, to provide the full range of possible states of interactions and average systems’ patterns trajectories. As causality is considered as non-linear predictability of diverse events of populations or communities, we believe OIF is the optimal model able to predict the largest divergence of trajectories due to the full consideration of ecosystem states via species probability distribution functions. Predictive causality is a convenient definition for any ecosystem, or data science problem more generally. However, for investigations of causality aiming to learn underlying physical processes of observed patterns, or for solving pressing issues of real complex ecosystems, a more in depth inquiry of complexity and dynamics (in relation to the target objectives), system learning and stakeholder collaboration are of paramount importance since data and models alone cannot reveal the full picture nor identify realistic and optimal solutions.

## 5 Materials and Methods

### 5.1 Ecosystems Models

#### 5.1.1 Bio-inspired Two Species Mathematical Model

In Sugihara et al. (2012) ^29^, a mathematical model was introduced to generate coupled nonlinear sequences for testing the CCM presented in that study. The model consists of two diffusively coupled logistic maps describing a simple bio-inspired dynamics without any external environmental effects on both species. It is analytically formulated as:

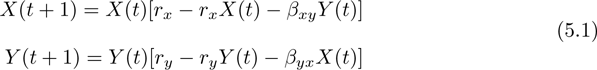

where *X* and *Y* are two random variables linked by factors *β_xy_* and *β_yx_* that establish the strength of their interactions. It gives possibility to estimate the “true” causality in an absolutely numerical sense and this model can be therefore indicated as *S*(*β_xy_, β_yx_*). If *β_xy_* is fixed as 0 and *β_yx_* is fixed as non-zero or varied free, that is, X causes Y, but not vice versa. If *β_xy_* and *β_yx_* are both non-zero, X causes Y and vice versa. These conditions generate two different kinds of coupling variables that respectively represent unidirectionally and bidirectionally interactive species-species systems described as the case 1 in Fig. 1. *r_x_* and *r_y_* are the intrinsic growth rates for each variable.

In this study, we focus on both unidirectionally and bidirectionally interactive species-species systems. For unidirectional coupling, *β_xy_* is fixed as 0, but *β_yx_* is free varied within the range of [0,1], leading to a simplified model as *S*(0*, β_yx_*). This unidirectional model *S*(0*, β_yx_*) is exploited to generate coupled sequences of two random variables X and Y where X affects Y, but not vice versa. For bidirectional coupling, we study two different conditions. On the one hand, we consider a system *S*(*β_xy_, β_yx_*) in which *β_xy_* is fixed as 0.2, 0.5 and 0.8, *β_xy_* is free varied within the range of [0,1]. This model can be indicated as a univariate model *S*(0.2*/*0.5*/*0.8*, β_yx_*) and is used to generate interdependent coupled time-series variables where the effect of X on Y changes over *β_yx_*, while the effect of Y on X is stablized due to the fixed non-zero *β_xy_* s. On the other hand, we consider a bivariate system *S*(*β_xy_, β_yx_*) in which both *β_xy_* and *β_yx_* are free varied. This model indicated as *S*(*β_xy_, β_yx_*) generates coupled time series of variables X and Y where X and Y randomly interact with each other. For all these conditions mentioned here, *S*(0*, β_yx_*), *S*(0.2*/*0.5*/*0.8*, β_yx_*) and *S*(*β_xy_, β_yx_*) are run under the conditions of which initial *x*(1) = 0.4, *y*(1) = 0.2 and intrinsic growth rates of variables *r_X_* and *r_Y_* respectively are 3.8, 3.5 as used in^29^. These time series of variables X and Y generated by the models for example can be time-series data of species abundance.

#### 5.1.2 Sardine-Anchovy-Temperature Ecosystem

As a real-world ecosystem case study, yearly time-series data of Pacific sardine landings, northern anchovy landings and sea surface temperature (SST) obtained at Scripps Pier and Newport Pier, California are used here. Sardines and anchovies seldom interact with each other because of geographical distribution. External environmental factors including SST affect sardines and anchovies, but not vice versa. It is a typical example of unidirectional causal relationship in real-world ecosystems and such tripartite relationship can be described as the case 2 in Fig. 1. Sugihara et al. (2012) ^29^ also studied this fishery ecosystem and successfully inferred the weak causal interactions between between sardines, anchovies and SST with the CCM method. We remake the experiments Sugihara et. al did, and apply our proposed OIF model to do the same work as well, thereby validate the OIF model by comparing results to those from CCM.

#### 5.1.3 Fish Community in the Maizuru Bay Ecosystem

Long-term time-series data counting the observations of the fish community collected along the coast of the Maizuru Fishery Research Station of Kyoto University ^37^ are used in the multispecies case study described as the case 3 in Fig. 1 for OIF model validation. Underwater direct visual censuses were conducted approximately once every two weeks from January 1, 2002 to April 2, 2014, totally generating 285 time points sequences during about 12 years long census. Dominant fish species (that is, with a total observation count was larger than 1,000) were considered, because rare species that were sporadically observed during most of the census term have many zero values, and yet there were not suitable for the time-series analysis. Information of rare species dynamics is typically very small within other species and their inclusion does not change results about the whole ecosystem interaction topology beyond just adding poorly or not interacting species as found in Ushio et al, (2018) ^37^. Therefore, only 14 dominant fish species and 1 jellyfish species were selected in this dataset. 1 Jellyfish species was selected in the dataset because this species was abundant in this area and was thought to have considerable influences on the community and ecosystem dynamics.

Accordingly, both OIF and CCM model are exploited to measure causal interactions among 14 dominant fish species and 1 Jellyfish setting up the dynamical complex multispecies system.

### 5.2 Interactions Inference Models

#### 5.2.1 Linear Correlation Model

The linear correlation between non-lagged random variables X and Y is given by:

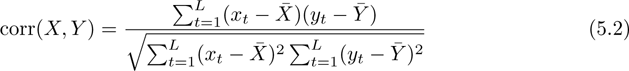

where 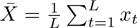 and 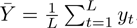. *L* is the length of time-series of X and Y.

#### 5.2.2 Convergent Cross Mapping Model

The principle of CCM model involves state space reconstruction from two variables and quantifies the potentially causal (asymmetrical) relationship between these variables using the method of nearest neighbor forecasting. Nearest neighbor forecasting method is an application of Takens’ Theorem called simplex projection. States of a system are reconstructed by applying successive time lags of time-series variable underlying the method of time lag embedding^29;38^. Interestingly, this method has been originally applied to describe the transition to turbulence of fluids ^69;70^.

In the case where X causes Y, Takens’ theorem indicates that there should exist a “causal” relationship between states of X and the contemporaneous states of Y. CCM quantifies this relationship using the simplex projection to predict time-series X from reconstructed Y. Specifically, a manifold *M_X_* (“reconstructed”, “shadow” or predictor manifold) is constructed from lags of variable X (i.e., *X*(*t − τ*) with the time lag *τ*) and used to estimate contemporaneous values of *Y* (*t*). *M_X_* is an approximation that will display convergence up to the level set by observational error and process noise. At convergence the approximated *Ŷ* (*t*)*|M_X_* will be close to *Y* (*t*). The relationship between *Y* (*t*) and *Y* (*t − τ*) is on the target manifold. To explore the opposite “causality” CCM explores the convergence of *X̂* (*t*)*|M_Y_* to *X*(*t*) where *M_Y_* is the predictor manifold. Thus, CCM determines how well local neighborhoods (defined by *E* + 1 points, that is the minimum number of points needed for a bounding simplex in an E-dimensional space) on the manifold *M_X_* correspond to local neighborhoods on *M_Y_*.

Pearson’s correlation coefficients *ρ* (originally defined as Eq. 5.2 considering non-time lagged variables) between predicted time series and observations of X (or Y) are calculated. The non-linear correlation coefficient is considered as the indicator of cross-mapping skill, that is the “causality” between species X and Y. The non-linear *ρ* is defined as:

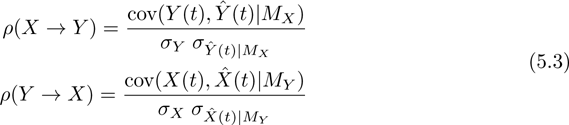

where cov and *σ* are the covariance and standard deviation. *X̂* (*t*)*|M_Y_* and *Ŷ* (*t*)*|M_X_* are the predicted values of *X*(*t*) and *Y* (*t*) considering the attractor manifolds of lagged Y and X. Considering the relationship between the calculated cross-mapping skill and the length of time series *L*, *ρ* increases with *L* until a convergent stable value. *ρ* is alway larger the longer *L* and that indicates causality according to Sugihara et al. (2012) ^29^. Typically, no less than 30 points in the time-series data should be used for CCM analyses ^71^. Further details about the use of CCM for this study are provided in Supplementary Information. CCM codes are available at https://github.com/HokudaiNexusLab/net-valid.

#### 5.2.3 Optimal Information Flow Model: TE Inference

OIF requires time-series data of variable biomarkers (e.g. abundance) as input, and produces a Transfer entropy (TE) matrix quantifying interactions between all pairs of variables. Therefore, the primary output of OIF is a functional interaction network for the ecosystem considered. In this paper we consider all inferred TEs without any redundant interaction removal as in Servadio and Convertino, (2018) ^49^. All OIF codes are available at https://github.com/HokudaiNexusLab/net-valid. TE is a non-parametric statistic in information theory that estimates the amount of information that a source variable contains about a destination variable considering destination’s current and historical states^40^. It measures how much directed (time-asymmetric) information transfers between two variables, giving an incentive to quantify the causal relationship between two variables with TE. Here it can be calculated as:

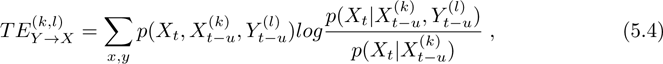

where, *X* and *Y* denote two random variables, *k* and *l* refer to the Markov orders of variables *X* and *Y*, implying that we need to at least consider *k* (*l*) time points of variable *X* (*Y*) in the past for the estimation in order to capture all relevant information in the past of *X* (*Y*). Here we assume that the time-series analysis here obeys a memoryless Markov process. Hence, parameters *k* and *l* are fixed as 1, that is to say, the next states of *X* and *Y* are only dependent on the current states and not on states in the past. *u* is the source-target time delay establishing lagged influence and is free varied. Servadio and Convertino (2018) ^49^ proposed a framework of optimal information networks (OIN) to select TEs that maximize the total entropy for inferred networks. Probability distribution functions associated to the network with maximum entropy are considered as the most predictive distributions fitting observations (where power-laws have the lowest entropy but highest uncertainty reduction when compared to other distributions). TE in Eq. 5.4 is estimated until the selected time *t* with incremental time series data whose minimum resolution is *g* (equal to 30 time points, that is 60 weeks) that defines the minimum number of points required for robust inference (see Supplementary Information section S1.3).

In this paper, OIF is improved by incorporating the well-documented JIDT toolkit ^42^ in computing TE. The JIDT toolkit provides users with multiple alternatives for pdf estimation including discrete, binned, Gaussian, Kraskov (KSG) and Kernel models that cover most data types (from uniform to power-law). Therefore, OIF with JIDT is more flexible in relation to diverse ecosystems and datasets.

JIDT-Kernel as model-free estimator calculates joint pdf by discretizing data into bins. A classical approach called Kernel estimation uses equal-width bins specified by a bin-width parameter. The joint pdf can be written as:

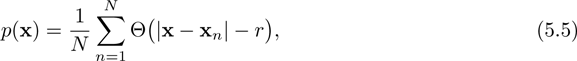

where Θ is a step kernel such that Θ(*x ≤* 0) = 1 otherwise equals zero, *|***x** *−* **x***_n_|* is the maximum distance between variable **x** and observation measurement **x***_n_* for *n* = 1, 2*, …, N*. The parameter *r* is the specified bin width. Note that the estimation is sensitive to the choice of parameter, and the number of bins can be different between variables. After calibration we selected the optimal bin-width as 0.25.

In this study we also investigate the optimal TE that provides a clearer detection and more accurate quantification for causal relationships between two variables by choosing an appropriate time delay *u* in a specified range. The choice of the optimal *u* within the range leads to the optimal TE model and resultant network inference while considering the minimum computational complexity. As shown in our previous work about microbiome ^39^, time delay *u* used to calculate TE between species is the one who minimizes the distance from one species to another in the inferred network.

The distance can be calculated by ^31^:

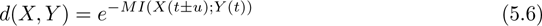

where MI is the mutual information of variables X and Y. MI of two random variables X and Y, is given by:

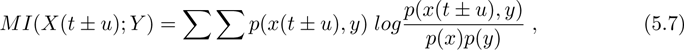

where *p*(*x*) and *p*(*y*) are the marginal distributions of random X and Y, and *p*(*x*(*t± u*)*, y*) is the joint probability distribution that is the pdf affected by the time-delay. The time delay *u* that is chosen is exactly the one that maximizes the MI of the two variables, because that is the one that minimizes the uncertainty. MIs is calculated using a range of time delays *u* (defining temporal entropy reduction parameters) and then the time delay corresponding to the maximum MI is selected before the calculation of TE. The choice of using the time delay *u* that maximizes MI is focusing on the highest predictability rather than the investigation of true causality. For example, two species may have a relatively low average interaction, except for a limited time period when the interaction is very high due to seasonal predation or sudden environmental disturbances (e.g., intense fishing or water pollution). Thus, by considering the maximum MI the focus is on extreme interactions (leading to the highest predictability) vs. average most likely mutual interactions, or more precisely it is focused on the magnitude of potential interactions rather than the interaction frequency. Yet, extreme accidental interactions are also captured by the choice of *MI*(*u* = *u_max_*), although very limited interactions may exist on average. This is equivalent to the approach of extended CCM^38^ for which the time delay that maximizes predictability (convergence) is chosen. The existence of a time delay explicitly takes into account delayed interaction effects, for example, due to delays in biomass conversion in case of predation; interestingly these model-inferred delays should be used when assessing predator-prey abundance scaling laws ^72^ (fingerprinting ecosystem metabolic function) whose exponent is affected. In addition, we also compare the results of the linear cross-correlation estimates and the non-linear MI estimates for species interactions. Mutual information is a distance between two probability distributions while correlation is a linear distance between two random variables. This is done to detect the causal relationships between variables in the non-linear mathematical predator-prey model (where the two variable X and Y can be also belong to the same species) also to detect the performance of linear vs. non-linear interaction inference models. On the contrary of the asymmetric TE (that measures directed interdependencies between variables), MI, as well as cross-correlation, provides a symmetric measure for inferring mutual interdependencies unable to identify the direction of potential causal interactions.

### 5.3 Predicted Ecosystem Biodiversity Patterns

Details about the calculations of taxonomic and effective *α*-diversity are contained in the Supplementary Information.

## Acknowledgments

M.C. and J.L. gratefully acknowledge the funding provided by the GI-CORE Global Station for Big-Data and Cybersecurity at Hokkaido University, Sapporo, JP. M.C. acknowledges the FY2020 SOUSEI Support Program and Award for Young Researchers awarded by the Executive Office for Research Strategy to the Top 20% scientists in terms of productivity and citations at Hokkaido University. M.C. also acknowledges the Microsoft AI for Earth computational resources and the NIH funded Big Data to Knowledge (BD2K) 2017 Innovation Lab “Quantitative Approaches to Biomedical Data Science Challenges in our Understanding of the Microbiome” managed by the BD2K Training Coordinating Center (TCC). Lastly, M.C. acknowledges the funding from the Key Technology Partnership program at the University of Technology Sydney “Biomimicry: Implementable Collective Swarm Dynamics”. We formally acknowledge reviewers’ effort whose interesting comments greatly improved the manuscript.

## Author Contribution

M.C. idealized the study, wrote the manuscript, and prepared all final figures. J.L. performed all calculations and created the preliminary figures.

