## Supplementary Information for "Inferring Ecosystem Networks as Information Flows"

**J Li**[a,b], **M Convertino**[a,b*]

[a] Nexus Group, Faculty and Graduate School of Information Science and Technology, Hokkaido University, Sapporo, JP

[b] GI-CORE Global Station for Big Data and Cybersecurity, Hokkaido University, Sapporo, JP

February 19, 2021

*Corresponding author:* * M. Convertino, 9 Chome, Kita 14, Nishi 9, Kita-ku, room 11-11, Graduate School of Information Science and Technology, Hokkaido University, Sapporo, Hokkaido, Japan, 060-0814, phone: +81 011-706-6491,

*Keywords*: complex networks, causality inference, ecosystems, transfer entropy, convergent cross mapping, dynamics

### S1 Supplementary Materials and Methods

#### S1.1 Data

All table and figures presented as Supplementary Information, expect for Fig. S1, are about the multispecies ecosystem in the Maizuru bay, JP.

#### S1.2 Convergent Cross Mapping

The number of time lags (affecting the size of time windows) is defined as embedding dimension $E$. The embedding dimension is the minimal dimension of the state space that allows the attractor reconstruction without topological ambiguity and able to represents all dynamic behaviors. $E \geq 2D + 1$ guarantees to reconstruct the original state space of a dynamical system (Theiler, 1990), where $D$ is the fractal dimension that, effectively, is a ratio providing a statistical index of complexity comparing how details in a pattern (strictly speaking, a fractal pattern, or "strange" attractors) change with the scale at which it is measured. for $E \geq D$ the reconstructed set will almost always have the same dimension as the attractor (Ruelle and Takens, 1971). The accuracy of reconstruction varies as a function of $E$. For instance, given a time-series variable $x(t)$, $E$-dimensional reconstruction is done by using successive lags of $x(t)$: $x(t)$, $x(t-\tau)$, $x(t-2\tau)$,..., $x(t-(E-1)\tau)$. Generally $\tau$ is fixed as 1 because most of the time series data are not over sampled and that allows to explore all sampled values without any assumption on the underlying dynamics. Because the performance of CCM is sensitive to the choice of the embedding dimension, $E$ should be carefully determined. In reality, selecting an optimal $E$ for state space reconstruction is a daunting task due to multiple factors including system complexity, size of time series and noise. In the rEDM package of CCM (Sugihara et al., 2012), convergent cross mapping is implemented in the *ccm* (https://rdrr.io/cran/rEDM/man/ccm.html) function in which the embedding dimension $E$ is regarded as a parameter that can be manually set by users. In this study, integers in a given range are used as embedding dimension $E$ to test the state space reconstruction (predictability) using univariate simplex projection (Sugihara and May, 1990). The $E$ that maximizes the predictability ensures to capture all relevant information of the system without including extraneous information in the past (or along other dimensions) (Ushio et al., 2018). In practice the embedding dimension (E) can be take equal to 3 ($D = 1$ because time series are considered), thus shadow manifolds are derived from 3 time-lagged coordinates (e.g., $[X(t), X(t-\tau), X(t-2\tau)] = M_X$ where $M_X$ is the manifold of X). The R package "rEDM" is available online https://rdrr.io/cran/rEDM/ through the Comprehensive R Archive Network (CRAN).

#### S1.3 Predicted Ecosystem Biodiversity Patterns

##### S1.3.1 Taxonomic and Effective $\alpha$-diversity

We consider the macroecological indicator $\alpha$-diversity (or taxonomic diversity) for the fish community in the Mairuzu Bay (Kyoto, Japan) to investigate whether the OIF model infers a "causal" network able to predict $\alpha$-diversity over time with high accuracy. Results are compared to the $\alpha$-diversity calculated from the CCM inferred network. The OIF and CCF $\alpha$-diversity are introduced as "effective" $\alpha$-diversity to consider the estimated interacting species rather than counting species independently of the interaction. Specifically, we base our analysis on taxonomic $\alpha$-diversity that is the most elementary definition of local biodiversity of a community. However, taxonomic $\alpha$ captures only one aspect of diversity that may not be sufficient especially for very uneven communities where species have very different abundance. Nonetheless, this does not affect our model intercomparison, since any model discussed here can be used for any diversity metric such as Shannon index and Simpson diversity.

$\alpha$-diversity is a concept in ecology that counts the number of species (biodiversity) observed at a local scale in space and time. In this multispecies case study, the local scale is a time-dependent measure for the whole community. The resolution at which $\alpha$ is assessed (i.e., the sampling interval of the time-series data) is two weeks. For a set of species $\mathbf{S} = \{S_1, S_2, ..., S_n\}$ whose abundance $\mathbf{X} = \{X_1, X_2, ..., X_n\}$ changes over time, $\alpha(t)$ can be calculated as:

$$\alpha(t) = \sum_{k=1}^{n} x_k(t)^0 , \tag{S1.1}$$

where $x_k(t)$ is the abundance of species $k$ at time point $t$.

An effective $\alpha$-diversity is also derived from the inferred species interaction networks using CCM and OIF models. The estimated effective $\alpha$-diversity (indicated as $\alpha_E(t)$, hereafter) is the number of nodes (species) in the inferred networks considering the minimum data length $l$ required for the inference. The estimated $\alpha$-diversity from CCM and OIF can be obtained as:

$$\alpha_E(g) = \sum_{i=1}^{n} k_i(g) \tag{S1.2}$$

where,

$$k_i(g) = \begin{cases} 0, & \text{for } \sum_{j=1}^{n}(|M_{i,j}(g)| + |M_{j,i}(g)|) = 0 \\ 1, & \text{for } \sum_{j=1}^{n}(|M_{i,j}(g)| + |M_{j,i}(g)|) \neq 0 \end{cases}, \tag{S1.3}$$

denotes the structural degree of all nodes (species) involved in the inferred networks for a time period $g = t - l$. $M(t)$ is the $n \times n$ interaction matrix from OIF or CCM model for each time period $g$.

The total number of time periods on which the inference of networks is performed (and $\alpha_E(g)$ is calculated) by both OIF and CCM is $G = \lfloor \frac{L-l}{\Delta t} \rfloor + 1$. $L$ is the total number of observations of abundance obtained every two weeks for each species, $l$ is the minimum number of observations set up to perform the inference of interactions (i.e., $l$=30 in this study supported by the evidence about the minimum length required to perform a robust inference of $\rho$ with CCM) for the whole time series $L$, and $\Delta t$ is the numerical inter-observation time (or time step) that corresponds to two weeks. $\lfloor \bullet \rfloor$ (where $\bullet = \frac{L-l}{\Delta t}$) rounds $G$ to the smaller integer. Note that $l$ is the maximum embedding dimension $E$ for CCM. In this multispecies case study, the length of raw time series is 285 and the time step is chosen as 1. Thus, G is equal to 256. CCM and OIF models leverage 256 shortened time series to estimate the potential causality between all possible pairs of fish species, leading to 256 dynamical networks. Note that the number of $\alpha(t)$ values is higher than the number of $\alpha_E(g)$ because of the need of a minimum data length of network inference models to infer species diversity.

##### S1.3.2 Simpson's Diversity and Shannon Indices

The Simpson's Diversity Index (SDI) has been calculated for comparison with $\alpha$-diversity. SDI is a macroecological indicator that gauges diversity differences in communities considering also species population abundance. The calculation of SDI is given by:

$$SDI(t) = 1 - \frac{\sum_{k=1}^{n}[x_k(t)(x_k(t)-1)]}{\sum_{k=1}^{n} x_k(t)\left(\sum_{k=1}^{n} x_k(t) - 1\right)};. \quad \text{(S1.4)}$$

The range of SDI is from 0 to 1 (1 is the total normalized diversity) in which high scores indicate high diversity and low scores indicate low diversity.

In light of the relationship between entropy and diversity introduced by Jost (2006), we also investigated the Shannon index that is defined as the sum of entropies of all species based on time-series abundance. This is the first component of the information balance equation as introduced in Li and Convertino (2019). The Shannon index is formulated as:

$$H_\alpha(t) = \sum_k H(x_k(t)) = -\sum_k p_k(t) \log p_k(t) \quad \text{(S1.5)}$$

where $p_k(t)$ is the probability to observe species $k$ at time point $t$; more specifically this probability is based on the abundance of each species at each time step. Probabilities are obtained from probability density function estimates via kernel density estimation (KDE). $H_\alpha(t)$ gives the uncertainty as diversity information index rather than the taxonomic diversity and describes how species are assembled together via their probabilistic nature. Distributions reflect dynamics of species instead of considering simple occurrence as a binary variable. We compare $H_\alpha(t)$ to taxonomic $\alpha(t)$ and SDI to recognize similarities and differences in biodiversity indicators (or patterns more generally). This emphasizes the differential sensitivity of interactions' importance for different indicators, whether one considers $\alpha(t)$ or SDI for instance. Different indicators, or patterns, reveal different information about the ecosystem analyzed.

### S2 Supplementary Results

Figs. S6B,C show time-varying estimated $\alpha$-diversity from CCM and OIF models after setting high thresholds for the correlation coefficient $\rho$ of CCM and TE of OIF, respectively. For illustrative purposes, the chosen threshold is the median value of the range of CCM $\rho$ ([-1,1]) and OIF's TE ([0,1]). Thus, the threshold for CCM correlation coefficient is 0 and for TE of OIF is 0.5. With these thresholds only species who have connected interactions beyond the threshold persist in inferred networks. In principle, in a deterministic sense the best threshold for capturing average $\alpha$-diversity and its fluctuations should be chosen for both models. However, based on a more realistic interpretation all species has an influence on others (as highlighted in Fig 7B,C), then the establishment of thresholds is more useful for exploring ranges of interdependencies and associated effective $\alpha$-diversity with respect to the average taxonomic diversity. In this sense it is interesting to explore the sensitivity of $\alpha$ (either from CCM or OIF) for low-, medium-, and high-interaction species that can highlight which set of species is more sensitive to external fluctuations for instance.

Fig. S6B shows only positive interactions for CCM while Fig. S6C shows highly interactive species for OIF. Both fluctuations of effective $\alpha$ present clear signatures of seasonality, that implies species with higher interactions (species 4-9) are more likely less sensitive to external factors, such as SST, because they follow environmental dynamics. Vice versa, species with low interactions such as species 1 and 2, are highly affected by external factors such as SST because they are more asynchronized with these environmental drivers and less able to rely on other species. From a preliminary analysis it is possible to see how species 1 and 2 increases and decreases their abundance for increasing SST whereas other species are less sensitive to variation in SST. From Fig. B,C it may be possible to conclude that CCM potentially overestimates diversity contributed by high interaction species but that is hard to judge since ranges of $\rho$ and TE are different and based on different analytics. Nonetheless, $\alpha_{TE>0.5}$ shows that effective diversity for high interactions is relatively small (average ∼5) as expected by the TE interaction matrix in Fig 7C. It is interesting to note how the trend of effective $\alpha_{TE>0.5}$ is increasing over time; a sign that species with high interaction are increasing in number while the overall average diversity is decreasing as highlighted by the taxonomic diversity and $\alpha_{CCM>0}$ to a certain extent. Then, OIF is attributing higher sensitivity to SST of low interaction species as previously mentioned, versus CCM that is predicting a broader sensitivity of all positively interacting species. In general, decreasing trends of real $\alpha$-diversity over time reflects negative influences of global temperature rise species assembly that can reduce the number of species in this region. CCM and seemingly OIF allow stakeholders to explore more specific questions such as which components of diversity are responsible for such diversity decline due to temperature sensitivity, or vice versa stable under change.

When considering the real fish ecosystem case study we compare different biodiversity indicators reflecting local diversity. Specifically, the taxonomic $\alpha$-diversity, Shannon entropy of multiple species (indicated as $H_\alpha(t)$) and Simpson's diversity index (SDI) at each time point were calculated (Fig. S7). Plots of taxonomic $\alpha$-diversity and SDI show obvious seasonal fluctuations. They fluctuate with increasing and decreasing temperature in an annual period; during winter months of intense fishing activity $\alpha$ goes down and during the summer it goes up. These results evidence that the number of species, their distribution and collective dynamics are directly affected by SST as well as other interdependent factors. $H_\alpha(t)$ captures more large scale trends while SDI captures fluctuations more.

When considering the whole time period, we observe that the global trend of $\alpha(t)$ presents a slight decrease and the fluctuations of SDI are higher. These are effects potentially caused by climate change or other anthropogenic changes such as highlighted in Comte and Olden (2017), Brander (2007) and Hein et al. (2014). Climate change and global warming negatively influences species diversity, species distributions and their collective dynamics by altering species' biological adaptation as well as inter- and intra-species interactions that determine ecosystem stability. By comparing $H_\alpha(t)$ with taxonomic $\alpha(t)$, these two indicators show similar seasonal fluctuations and average trajectory over time. These seasonal fluctuations in $H_\alpha(t)$ show that fish species are more evenly distributed during periods with lower temperature due to the fact $H_\alpha(t)$ is higher. In addition, $H_\alpha(t)$ shows the same decreasing global trend in the whole period. Also, it presents similar high fluctuations in a yearly time scale which can be also observed in the plot of SDI. These results imply that in this case study the information-theoretic diversity (Shannon index) provides more information and a better performance in reflecting changes in species diversity, which can be applied for the prediction of macroecological patterns.

### S3 Supplementary Discussion

#### S3.1 "Causality" Inferential Methods: CCM and OIF

*Correlation* allows one to quantify the trend for linear systems but cannot be used to predict changes in biomarker values (e.g. abundance) of individuals and populations. Correlations between variables may signify the existence of non-linearity between of the variable considered and others. The fact that correlation between species abundances is somewhat proportional to TE implies that both measures provide a good estimate about the average magnitude of predictable interactions for multiple species in populations. It is worth to underline that correlation measures an average trend but no more information beyond that. $\rho$ allows one to quantify the non-linear trend and can be used to determine values of biomarkers such as population abundance. This is because $\rho$ is a metric that can be plugged in a set of non-linear differential equations predicting species abundance. *TE* allows one to quantify the change in pdfs and the likely range of biomarkers (species abundance in this context) associated to the observed changes in other variables. Additionally TE gives the information of the time delay between species. Only a recent modification in the CCM model is able to provide the temporal delay for $\rho$ (Ye et al., 2015) informing about the characteristic temporal scale of dependency between variables.

Causality principles and models, capturing only pieces of the causality problem that have been invoked in the past are the following.

- *Causal Sufficiency.* This is about the set of measured variables that include all of the common causes of pairs. This is obviously not always occurring and hard to even identify whether all variables that are potentially causal are observed. Additionally, this in an element that makes true causality inference difficult as well as not an element that impedes to develop an accurate predictive model.

- *Principle of independent mechanisms.* An example is prediction invariance, that is if by adding one variable predictions change then the added variable is not causal for the predicted outcome. This is obviously false because any prediction is dependent on the choice of the model and on spatio-temporal scales that defines the structure of the model. It is certainly true that causal models (defined as the ones that contain approximately all causal factors) are likely the ones that provide scale-invariant predictions but not necessarily since these models may vary in complexity and variable independency may vary.

- *Reichenbach's common cause principle.* If variables are dependent then they are either causal to each other (in either direction) or driven by a common driver. This is certainly a false statement because dependency does not imply true causality and dependency may be driven by different dynamics responding to different external stimuli (also when these stimulate change mildly but rapidly). An example is regular and scale-free dynamics that are likely highly dependent in a TE sense (where TE from scale-free to regular dynamics is high but low vice versa due to the fact scale-free dynamics can predict regular dynamics, such as in the case of epidemic-endemic dynamics) and driven by different environmental drivers. Models that just look into statistical dependence typically do not include directionality and asymmetry in causality due to space-time factors.

- *Odd ratio.* This is about the ratio of probabilities with and without a selected variable considered to be causal considering one variable at a time like one factor at a time sensitivity analyses. These models are based on ratios calculated on single probability values of deterministic and discrete presence/absence scenarios of causal factors. These models do not include directionally and asymmetry in causality but more importantly do not recognize the probabilistic character of complex system dynamics where the non-linear systemic interplay of variables determines the causal outcome of interest.

#### S3.2 Transfer Entropy and Predictability

TE is directly proportional to the divergence and asynchronicity of the variables' pdfs; the higher the difference in pdfs at the same and different time points the higher TE. In a predictability viewpoint, high TE implies high ability to predict the mutual change of variables and potential high interdependence between these variables. High TE is for positive interactions in a predictive sense (positive because one variable allows to predict another). Positive interaction is likely showing divergent and asynchronous behavior in species where asynchrony is more important than divergence (based on the calculation of TE); yet, for instance abundance is changing in opposite directions with time delays. Thus, a high correlation coefficient does not imply convergence (synchronous similarity between pdfs) because correlation works well only for linear systems. Therefore any conclusion based on correlation only for highly non-linear systems must be avoided. Small TE is for small interactions and low predictability; in this situation the dynamics is much more regular. High TE is for high interactions and high predictability; in this situation the dynamics is much more scale-free and species display opposite behavior to the same external stimuli. If two species abundances go into the same direction then TE is likely small. Biologically speaking the magnitude of TE is much more about interaction magnitude than frequency and then TE is focused on critical interactions with low frequency. Theoretically, negative values of $\rho$ mean that species mutual predictability is zero. These values are often set up to zero. Computationally, the negative values can be interpreted as misinformation, i.e. the information that predicts opposite trajectories with respect the observed ones. It is interesting to see that very small values of TE (corresponding to low predictability) are associated to negative values of $\rho$.

It is well known that MI is a quantity that measures how much information is shared between two random variables on average. In other words, it quantifies the reduction in uncertainty of one variable (entropy reduction) given the knowledge of another for a certain time delay (Villaverde et al., 2014). Zero mutual information between two random variables means the variables are independent. In the bio-inspired mathematical model formulated as in Eq. 1, current $X(t)$ and $Y(t)$ are used to respectively generate the next time point of themselves $X(t+1)$ and $Y(t+1)$. The length of historical values used to generate the coupling time-series data is 1. The amount of information that $X(t)$ contains about $Y(t+1)$ and that $Y(t)$ contains about $X(t+1)$ (MI) most likely is greater. Therefore, we choose 1 as the time delay $u$ used for TE calculations in this mathematical case study. In the sardine-anchovy-temperature system, no significant causal association between sardine and anchovy landings is observed (Lindegren et al., 2013), yet sea surface temperature (SST) affects all life including sardines and anchovies in the ocean. Sardines flourish when waters are warmer than average, while anchovies become the dominant small fishes in the Pacific when waters are cooler (Chavez et al., 2003). However, not only due to the weak to moderate effects of SST on sardines and anchovies, but also because both sardines and anchovies can live through a very wide temperature range, SST might have a long time delay to greatly affect the lives of sardines and anchovies in the ocean (Lluch-Belda et al., 1991). Accordingly, in order to accurately investigate the causal interactions between SST and sardines and anchovies, we use all integers in the range of [1,20] as multiple time delays for TE calculations in this case study. In the case of fish community, 14 fish species and 1 jellyfish are involved in the census, resulting in a complex ecosystem composed of large number of interspecific interdependencies among these species. Choosing a specific time delay for each TE calculation is computation-intensive because of highly nonlinear dynamics. Here we first acquire all time delays for TE calculations based on the proposed OIF model and then estimate the probability density function of these selected time delays. To simplify the calculation, we choose one time delay with a higher probability that approximately makes most MIs relatively greater according to the probability density function to calculate all TEs. In this fish community case study, the time delay can be 1 considering the probability distribution of all time delays shown in Fig. S8. In a word, in this study we don't choose the specific optimal condition for each TE estimation, but just consider the global optimum for the OIF model.

##### S3.3 Transfer Entropy, Ecosystem Stability and Shifts

In between species competition is good and we refer to this as a positive interaction also in a predictive sense because it helps in predicting one species from another. This corresponds to high TEs. Li and Convertino (2019) observed that microbiome unhealthy states are characterized by excessive divergence or competition characterize by an increase in TEs. In non-linear systems it also important to consider the least interacting species as well because these may be responsible for the maintenance of certain ecosystem states. For example in Li and Convertino (2019) the most detrimental bacteria in the microbiome were observed as the most abundant species that are characterized by the lowest Outgoing Transfer Entropy (OTE). Too much cooperation (or convergence) and disorganization can also lead to instability. Cooperation implies low predictability in terms of TE due to the fact that no major variations in one species are observed with respect to changes into the others. This occurs when many small value of TEs are observed which implies diffused disorganized interactions with high frequency and low magnitude. Criticality is the optimal stable situation in which there is a good organized balance between competition and cooperation although multiple stable states may exist associated to different interaction network topologies. It is indeed about both interactions magnitude and distribution. In critical ecosystems, where collective abundance is power-law distributed, divergent species are typically exponentially distributed (and yet more sensitive to environmental changes) in term while convergent species are power-law distributed. Relative stability is typically implying predictability. The ideas presented may caution modelers about the fact that unpredictability may be caused by either lack of monitoring or true causal independence of one species given the others. Therefore speculations of inferred interactions in the biological realm should be done with care and a safer choice is to focus on predictability rather than causality. We believe it is absolutely crucial to talk about TE in terms of systems predictability rather than causality, or equivalently to interpret causality as (information based) predictability rather than true biological causality. It is not just about methodological issues that render causality assessment difficult but also monitoring and data type considerations (in relation to inferential modes used) that contribute to the complexity of the causality inference problem.

### Supplementary Table Captions

**Table S1. Maizuru fish community species information**. Species ID considering the Maizuru dataset, scientific and common name, categorization in terms of fish stock, location endemicity (native/invasive), and reported IUCN conservation status (up to June 2020).

**Table S2. Comparison of inferred average interaction by CCM and OIF models**. Number referring to selected species interactions as in Ushio et al. (2018) for establishing the directional interaction from source to target species, CCM $\langle \rho \rangle$, OIF $\langle TE \rangle$ and associated time delay of TE. The estimated time delay associated to $\langle \rho \rangle$ is not reported.

**Table S3. Entropic and abundance characterization of Maizuru bay species considering the OIF model.** Species ID considering the Maizuru dataset, Shannon entropy based on pdf of abundance, Outgoing and Incoming TE where TE is from the OIF model, and mean and standard deviation of species abundance.

### Supplementary Figure Captions

**Figure S1. Relationship between suboptimal $\rho$ and TE**. Suboptimal $\rho$-s and TEs for all selected library length $L$ of time series and the time delay $u$ between variables (sardine-anchovy, sardine and SST, anchovy and SST from top to bottom), respectively, are shown.

**Figure S2. Interspecies abundance pattern**. Abundance-abundance patterns of all species in the Maizuru bay independent of time. The higher the correlation coefficient the higher the divergence and asynchronicity between abundance time series, and the higher TE (see Fig. 7 for species from 4 to 9). "Mirage" correlation between abundance of species (without considering the delay between autocorrelated values) implies non-linearity and potentially strong causality/physical interaction as demonstrated in Fig. 4 by the mathematical model results. Vice-versa, lack of correlation or low correlation implies linearity into the dynamics and potentially low causality/physical interaction. TE is advantageous because it is asymmetrical while interspecies correlation is symmetrical, yet not allowing one to capture the directional interaction between species.

**Figure S3. Time-varying interspecies interactions via OIF and CCM model for the Maizuru bay fish community.** Interspecies interactions for the 14 pairs of fish species listed in Table S2 quantified via (A) OIF and (B) CCM models. The y-axis indicates the time period from 2002 to 2014 over which the abundance of species was sampled every two weeks.

**Figure S4. Mean interaction strength and dominant eigenvalue from OIF and CCM interaction matrix.** (A) Average of all species-species interactions over time; (OIF and CCM model estimates in red and blue). (B) Atemporal relationship between mean interactions from OIF and CCM models. (C) Dominant eigenvalue corresponding to the highest frequency in interactions fluctuations (OIF and CCM model estimates in red and blue); the real part of the dominant eigenvalue of the interaction matrix at each time point is reported and represents a potential dynamical stability. (D) Atemporal relationship between dominant eigenvalue from OIF and CCM models. For (B) and (D) the slope $k$ is the liner regression coefficient.

**Figure S5. Distribution of pairwise species interactions from CCM and OIF**. (A) pdf of TE for all species interactions in the period 2002-2014. (B) pdf of $\rho$ for all species interactions in the period 2002-2014. (C) TE and CCM calculated for each species pair $ij$ considering each time period where abundance time series are updated every two weeks in the period 2002-2014. The number of estimated TEs and $\rho$ (i.e., 3584) is given by the number of observations $N$ (i.e., 256) multiplied by the number of species pairs (i.e., 14).

**Figure S6. Predicted $\alpha$-diversity via CCM and OIF versus taxonomic diversity.** (A) "Real" taxonomic $\alpha$-diversity (green line), and inferred temporal $\alpha$-diversity from CCM and OIF (red lines) without setting any threshold on the magnitude of species interactions ($\rho$ and TE). (B) Inferred $\alpha$-diversity after setting the threshold to zero for CCM $\rho$ (see Fig. S5 for pdf of $\rho$ where $\rho$ can be negative). (C) Inferred $\alpha$-diversity from OIF after setting the threshold to 0.5 for TE (see Fig. S5 for pdf of TE). TEs are obtained from JIDT using a time delay $u = 1$ that corresponds to 2 weeks. Sample data are provided every two weeks for 12 years (see Fig. 7).

**Figure S7. Biodiversity indicators over time for the Maizuru bay fish community.** (A) Taxonomic $\alpha$-diversity as count of diverse species. (B) Shannon diversity index $H_\alpha(t)$ based on the pdf of population abundance of species at each time step (Eq. 2.8). (C) Simpson's diversity index (SDI) over time that measures diversity difference based on population abundance at adjacent time steps (Eq. 2.7). $H_\alpha(t)$ is capturing more the trend of biodiversity that is in this case decaying over time in terms of abundance but increasing in regularity because of the lower entropy. SDI is more related to fluctuations whose periodicity is getting more stable in this ecosystem and observable in the autocorrelation of $\alpha$.

**Figure S8. Network entropy dependent on the TE threshold.** Network entropy dependent on the pairwise information flow (TE) between species. Network entropy is defined as the sum of Shannon entropies of all species (considering abundance) and TEs of all pairwise species interactions as in Li and Convertino (2019).

**Figure S9. Pdf of time delay for TE**. $u$ is the time delay that minimizes the statistical distance defined as $\exp^{-MI(X,Y)}$, where MI is the Mutual Information between species X and Y as in Li and Convertino (2019). The elementary unit or resolution of time delay $u = 1$

corresponds to the species sampling of two weeks.

| Species | Fish Stock | Native/Invasive | Conservation Status |
|---|---|---|---|
| 1. *Aurelia aurita* (Moon jellyfish) | Yes | Native | NE |
| 2. *Engraulis japonicus* (Japanese anchovy) | Yes | Native | LC |
| 3. *Plotosus lineatus japonicus* (Sea catfish) | No | Invasive | NE |
| 4. *Sebastes inermis* (Black snapper) | Yes | Native | LC |
| 5. *Trachurus japonicus* (Horse mackerel) | Yes | Native | NT |
| 6. *Girella punctata* (Blackeye seabream) | Yes | Native | NE |
| 7. *Pseudolabrus sieboldi* (Wrasse) | Yes | Native | LC |
| 8. *Halichoeres poecilopterus* (Rainbow wrasse) | Yes | Native | LC |
| 9. *Halichoeres tenuispinnis* (Chinese wrasse) | No | Invasive | LC |
| 10. *Chaenogobius gulosus* (Goby) | No | Native | NE |
| 11. *Pterogobius zonoleucus* (Blue/Yellow striped Goby) | No | Native | LC |
| 12. *Tridentiger trigonocephalus* (Chameleon goby) | No | Native | NE |
| 13. *Siganus fuscescens* (Rabbitfish) | Yes | Invasive | LC |
| 14. *Sphyraena pinguis* (Red barracuda) | Yes | Native | NE |
| 15. *Rudarius ercodes* (Pigmy filefish) | No | Invasive | LC |

Table S1:

| **Number** | **Source** | **Target** | $\langle \rho \rangle$ | $\langle TE \rangle$ | **u(TE)** |
|---|---|---|---|---|---|
| 1 | S. pinguis | T. japonicus | -0.2451 | 0.0886 | 0 |
| 2 | T. japonicus | S. pinguis | 0.0701 | 0.1029 | 1 |
| 3 | T. japonicus | Aurelia a. | 0.1416 | 0.4881 | 0 |
| 4 | H. tenuispinis | P. poecilepterus | 0.5860 | 0.5781 | 0 |
| 5 | P. poecilepterus | S. cheni | 0.0718 | 0.6516 | 0 |
| 6 | P. l. japonicus | P. sieboldi | -0.1407 | 0.1387 | 7 |
| 7 | T. trigonocephalus | C. gulosus | 0.0241 | 0.1393 | 2 |
| 8 | S. fuscescens | P. poecilepterus | -0.3727 | 0.1469 | 0 |
| 9 | G. punctata | P. zonoleucus | 0.1606 | 0.3199 | 0 |
| 10 | P. l. japonicus | T. trigonocephalus | -0.2120 | 0.1253 | 9 |
| 11 | R. ercodes | T. japonicus | 0.1606 | 0.4739 | 0 |
| 12 | P. zonoleucus | R. ercodes | 0.0210 | 0.3427 | 0 |
| 13 | P. zonoleucus | C. gulosus | -0.0124 | 0.0990 | 0 |
| 14 | P. zonoleucus | P. sieboldi | 0.0828 | 0.3734 | 0 |

Table S2:

| Species | Shannon Entropy | OTE | ITE | Mean | Std |
|---|---|---|---|---|---|
| 1. Aurelia aurita | 2.5031 | 6.4767 | 9.612 | 23.826 | 126.05 |
| 2. Engraulis japonicus | 0.5068 | 8.1583 | 3.4124 | 68.056 | 379.3 |
| 3. Plotosus lineatus japonicus | 0.5472 | 7.0584 | 3.1744 | 28.253 | 133.03 |
| 4. Sebastes inermis | 0.3261 | 6.16 | 10.524 | 31.874 | 50.512 |
| 5. Trachurus japonicus | 0.9996 | 6.3931 | 8.029 | 174.94 | 258.7 |
| 6. Girella punctata | 0.9914 | 6.0511 | 7.7665 | 14.863 | 29.911 |
| 7. Pseudolabrus sieboldi | 0.5570 | 6.2727 | 10.308 | 7.3333 | 6.9701 |
| 8. Halichoeres poecilopterus | 0.9803 | 6.1276 | 6.978 | 7.5649 | 11.755 |
| 9. Halichoeres tenuispinnis | 0.9903 | 6.6597 | 5.7889 | 17.575 | 33.523 |
| 10. Chaenogobius gulosus | 0.4852 | 8.0477 | 2.5391 | 8.9386 | 51.713 |
| 11. Pterogobius zonoleucus | 0.9379 | 6.2296 | 6.8386 | 18.542 | 77.554 |
| 12. Tridentiger trigonocephalus | 0.2022 | 6.9856 | 11.227 | 31.458 | 43.289 |
| 13. Siganus fuscescens | 0.4855 | 7.0709 | 3.7683 | 4.6456 | 25.866 |
| 14. Sphyraena pinguis | 0.3922 | 8.1649 | 2.3686 | 8.7614 | 50.779 |
| 15. Rudarius ercodes | 0.5852 | 6.0053 | 9.527 | 12.142 | 33.463 |

Table S3:

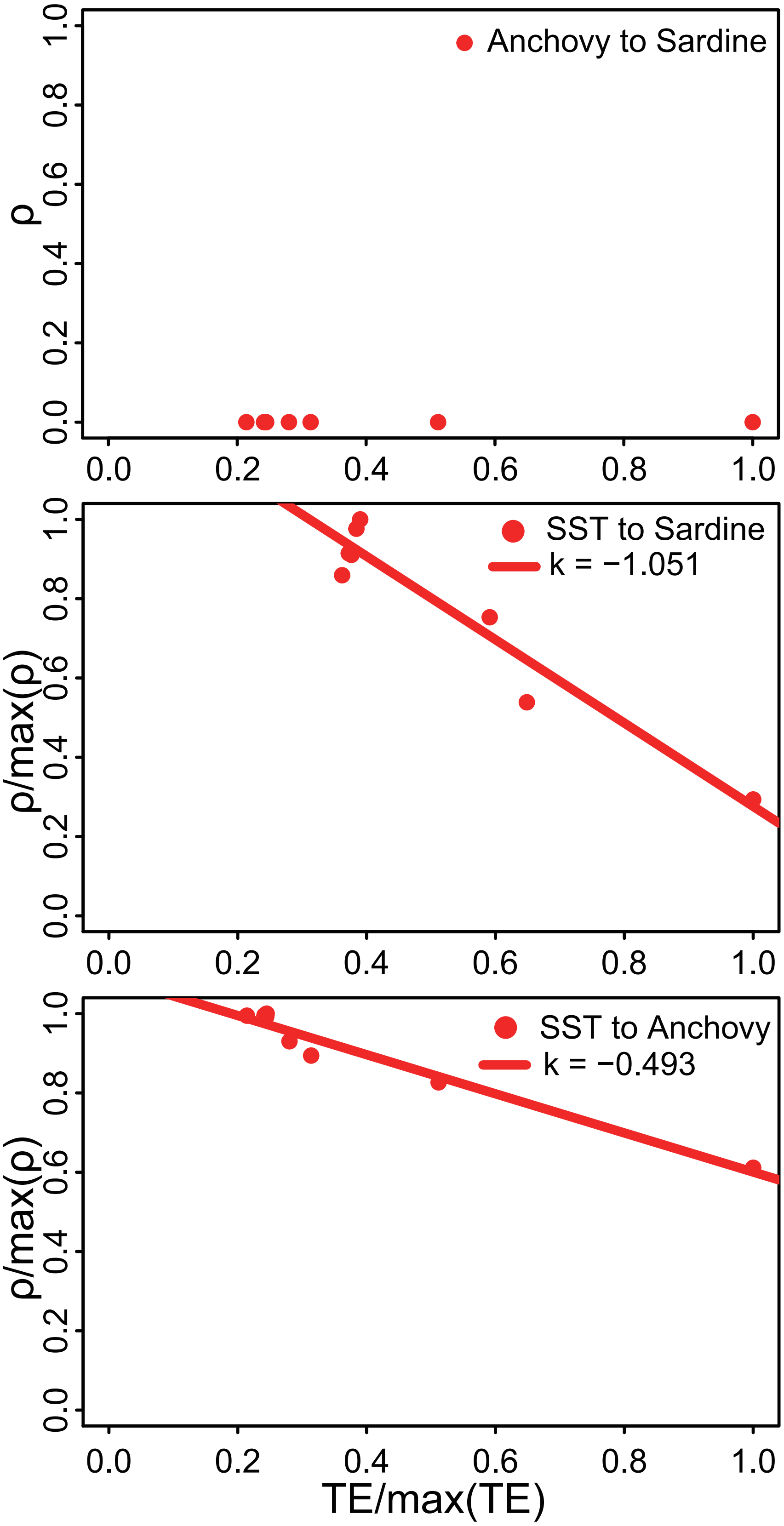
Anchovy to Sardine
SST to Sardine
k = −1.051
SST to Anchovy
k = −0.493
ρ
ρ/max(ρ)
ρ/max(ρ)
TE/max(TE)
0.0
0.2
0.4
0.6
0.8
1.0

Figure S1:

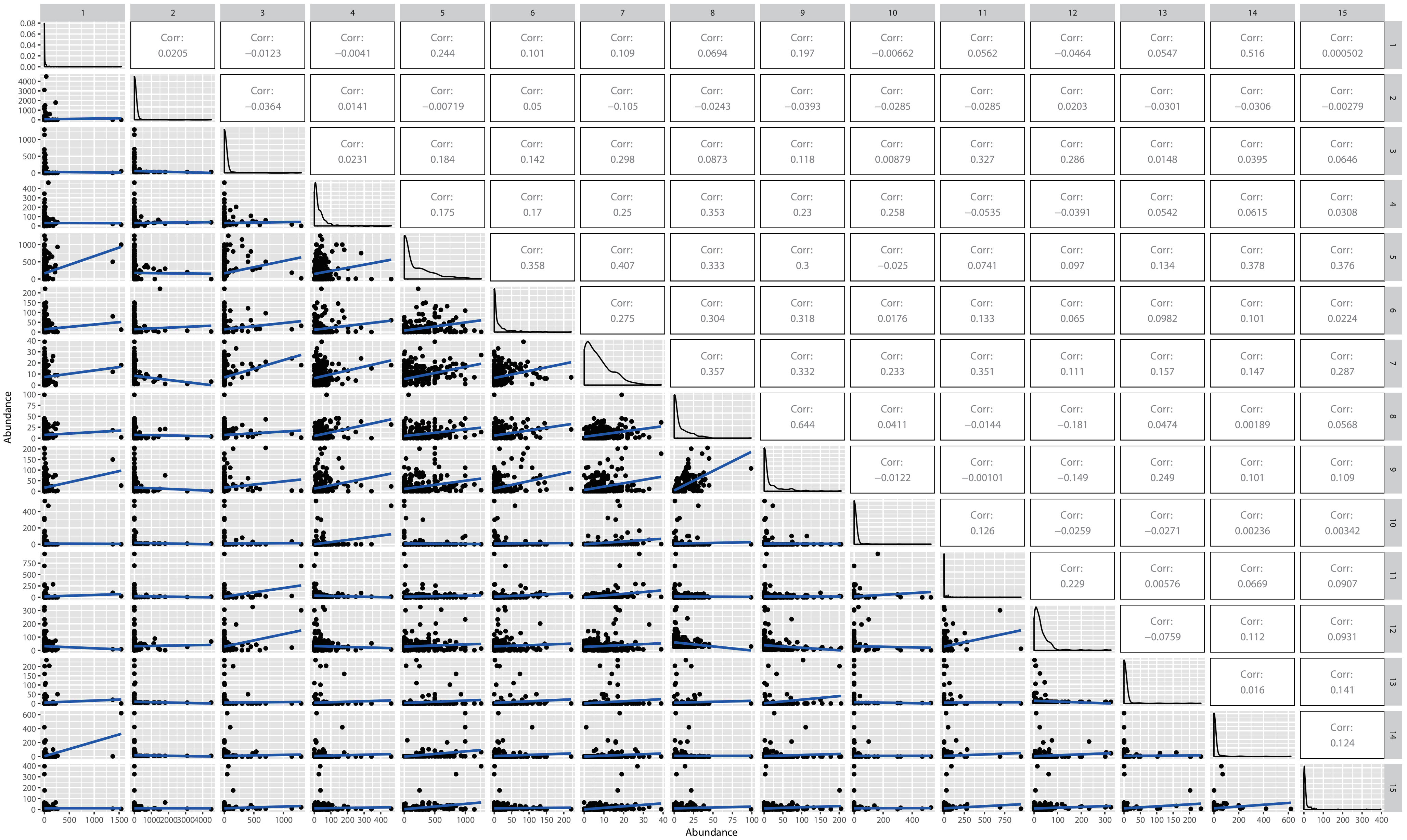
Corr: 0.0205
Corr: −0.0123
Corr: −0.0041
Corr: 0.244
Corr: 0.101
Corr: 0.109
Corr: 0.0694
Corr: 0.197
Corr: −0.00662
Corr: 0.0562
Corr: −0.0464
Corr: 0.0547
Corr: 0.516
Corr: 0.000502
Corr: −0.0364
Corr: 0.0141
Corr: −0.00719
Corr: 0.05
Corr: −0.105
Corr: −0.0243
Corr: −0.0393
Corr: −0.0285
Corr: −0.0285
Corr: 0.0203
Corr: −0.0301
Corr: −0.0306
Corr: −0.00279
Corr: 0.0231
Corr: 0.184
Corr: 0.142
Corr: 0.298
Corr: 0.0873
Corr: 0.118
Corr: 0.00879
Corr: 0.327
Corr: 0.286
Corr: 0.0148
Corr: 0.0395
Corr: 0.0646
Corr: 0.175
Corr: 0.17
Corr: 0.25
Corr: 0.353
Corr: 0.23
Corr: 0.258
Corr: −0.0535
Corr: −0.0391
Corr: 0.0542
Corr: 0.0615
Corr: 0.0308
Corr: 0.358
Corr: 0.407
Corr: 0.333
Corr: 0.3
Corr: −0.025
Corr: 0.0741
Corr: 0.097
Corr: 0.134
Corr: 0.378
Corr: 0.376
Corr: 0.275
Corr: 0.304
Corr: 0.318
Corr: 0.0176
Corr: 0.133
Corr: 0.065
Corr: 0.0982
Corr: 0.101
Corr: 0.0224
Corr: 0.357
Corr: 0.332
Corr: 0.233
Corr: 0.351
Corr: 0.111
Corr: 0.157
Corr: 0.147
Corr: 0.287
Corr: 0.644
Corr: 0.0411
Corr: −0.0144
Corr: −0.181
Corr: 0.0474
Corr: 0.00189
Corr: 0.0568
Corr: −0.0122
Corr: −0.00101
Corr: −0.149
Corr: 0.249
Corr: 0.101
Corr: 0.109
Corr: 0.126
Corr: −0.0259
Corr: −0.0271
Corr: 0.00236
Corr: 0.00342
Corr: 0.229
Corr: 0.00576
Corr: 0.0669
Corr: 0.0907
Corr: −0.0759
Corr: 0.112
Corr: 0.0931
Corr: 0.016
Corr: 0.141
Corr: 0.124
Abundance
Abundance

Figure S2:

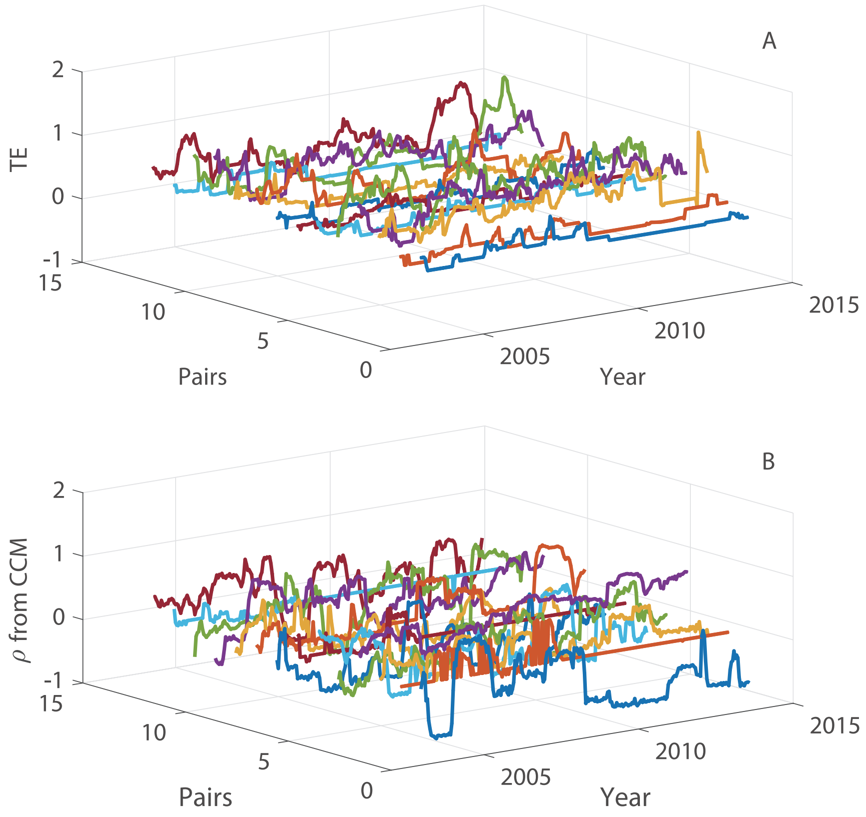
A
2
1
0
-1
TE
15
10
5
0
Pairs
2005
2010
2015
Year
B
2
1
0
-1
ρ from CCM
15
10
5
0
Pairs
2005
2010
2015
Year

Figure S3:

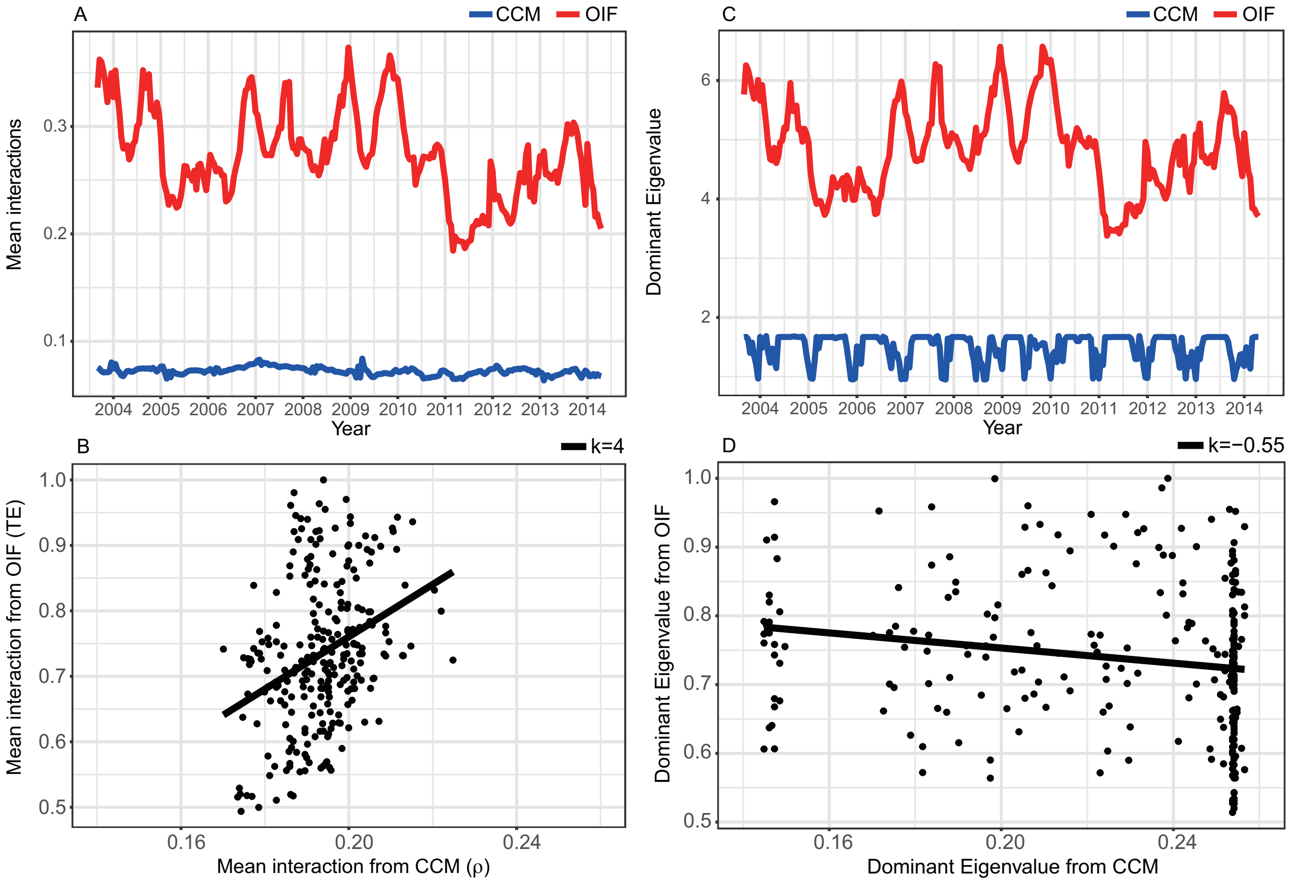
A
CCM
OIF
Mean interactions
0.3
0.2
0.1
2004 2005 2006 2007 2008 2009 2010 2011 2012 2013 2014
Year
C
CCM
OIF
Dominant Eigenvalue
6
4
2
2004 2005 2006 2007 2008 2009 2010 2011 2012 2013 2014
Year
B
k=4
Mean interaction from OIF (TE)
1.0
0.9
0.8
0.7
0.6
0.5
0.16
0.20
0.24
Mean interaction from CCM (ρ)
D
k=−0.55
Dominant Eigenvalue from OIF
1.0
0.9
0.8
0.7
0.6
0.5
0.16
0.20
0.24
Dominant Eigenvalue from CCM

Figure S4:

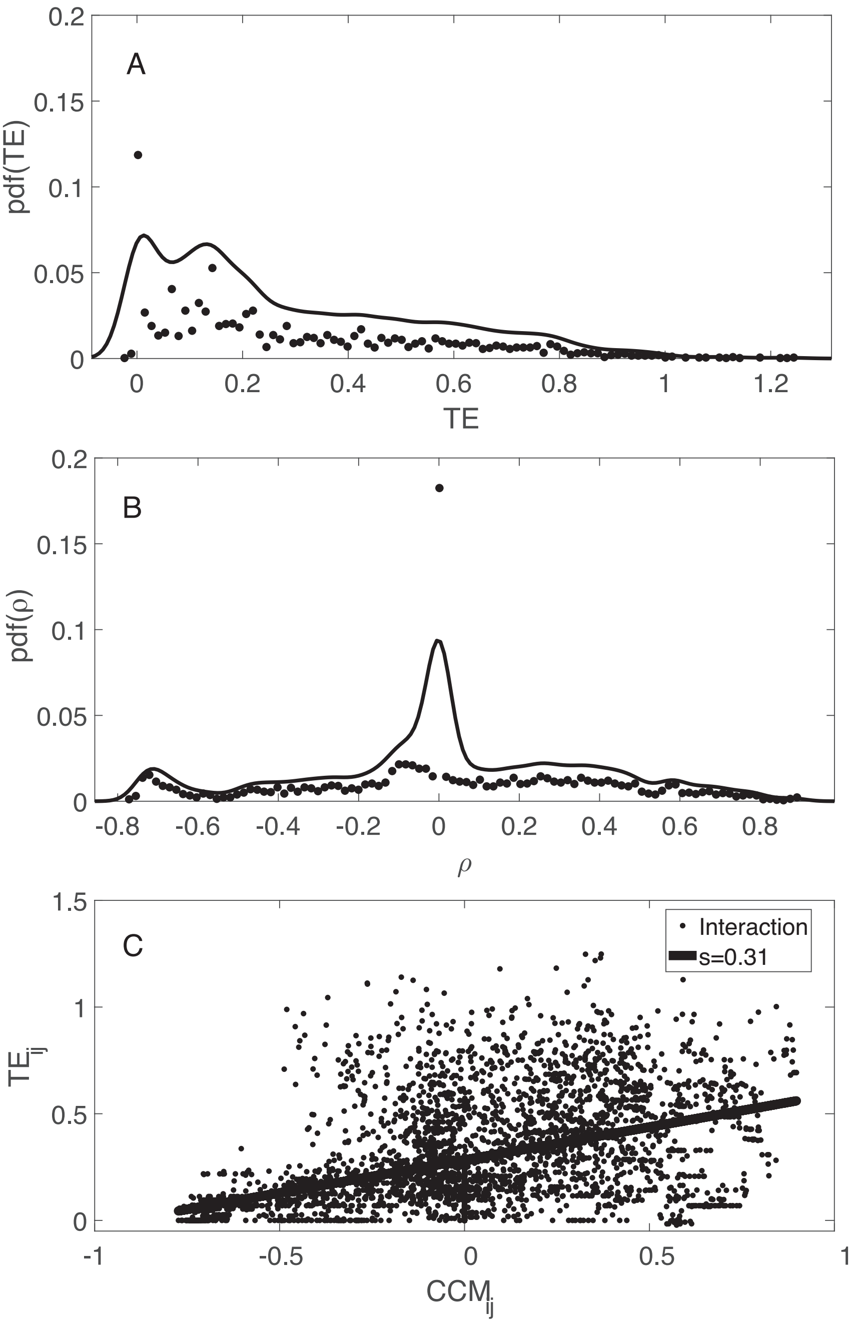


Figure S5:

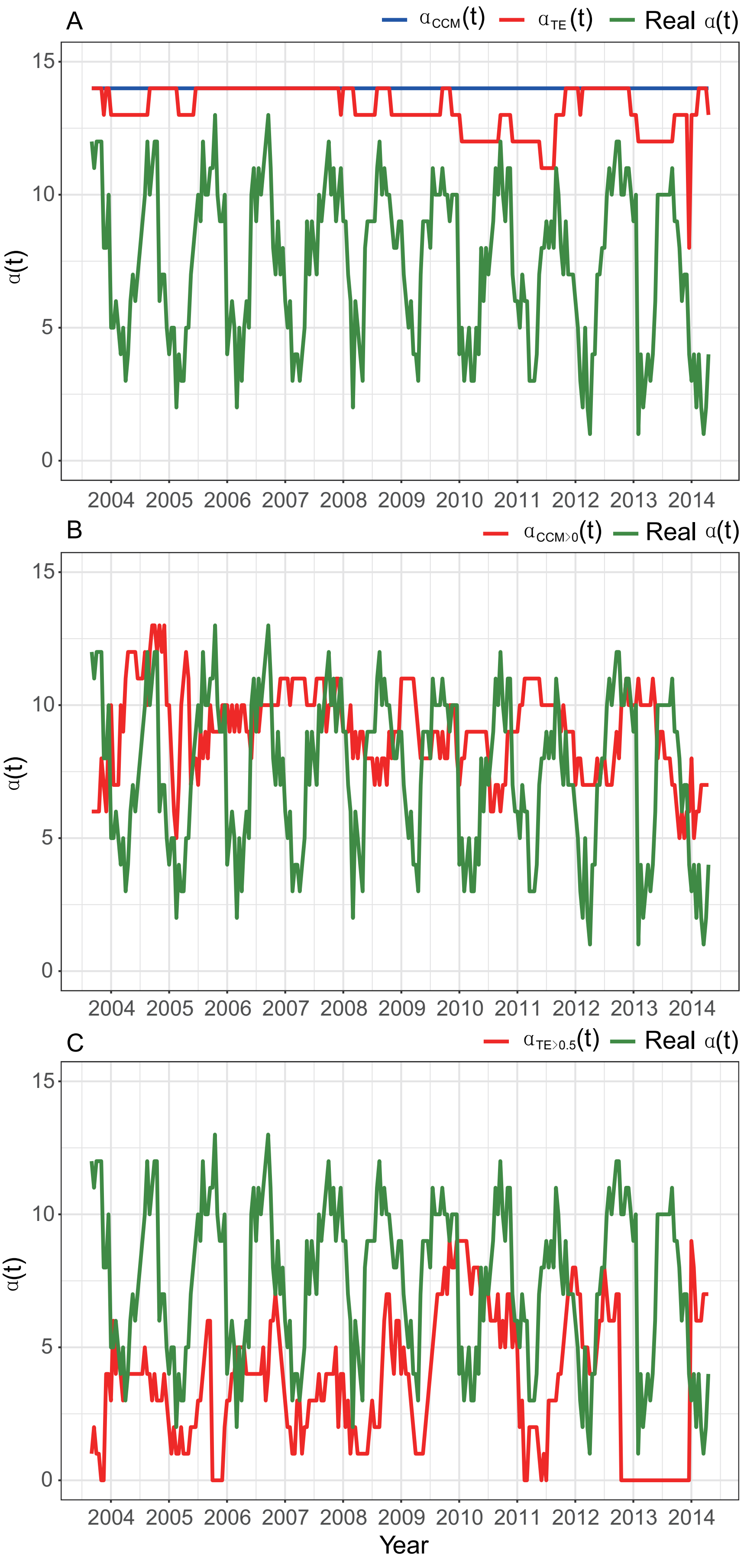
A
αCCM(t)
αTE(t)
Real α(t)
B
αCCM>0(t)
Real α(t)
C
αTE>0.5(t)
Real α(t)
α(t)
0
5
10
15
2004 2005 2006 2007 2008 2009 2010 2011 2012 2013 2014
Year

Figure S6:

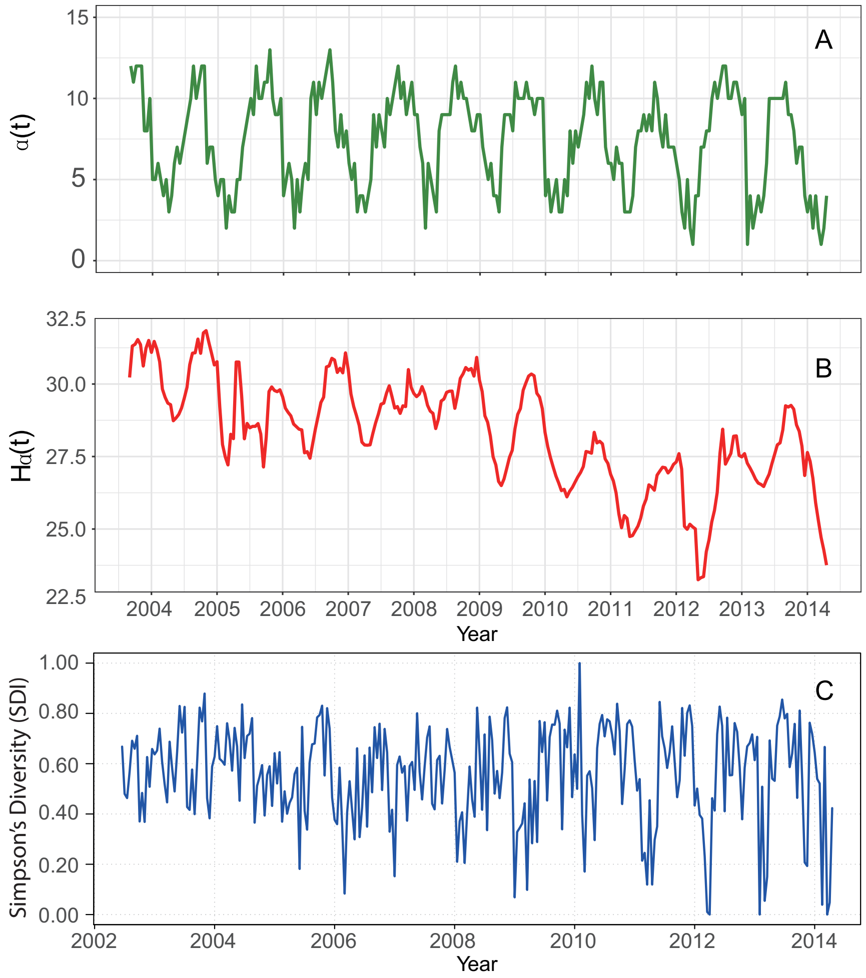


Figure S7:

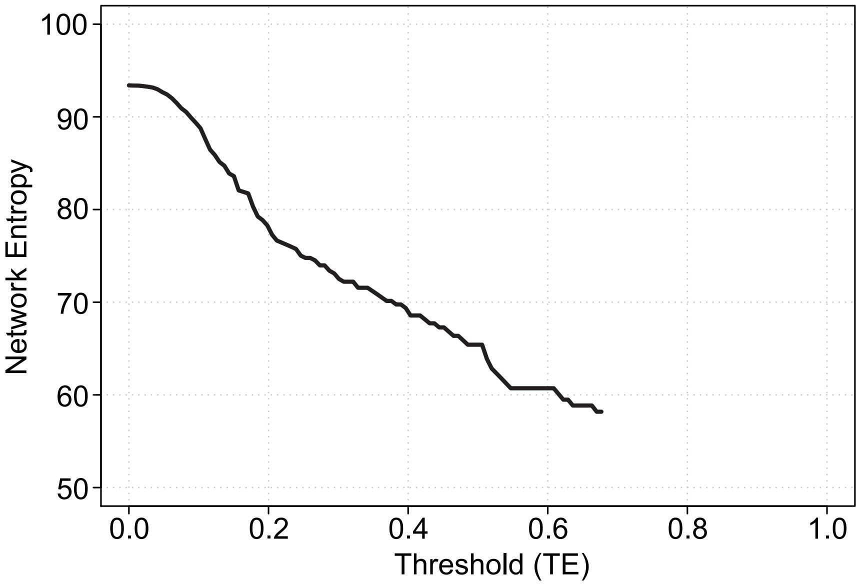
Network Entropy
100
90
80
70
60
50
0.0
0.2
0.4
0.6
0.8
1.0
Threshold (TE)

Figure S8:

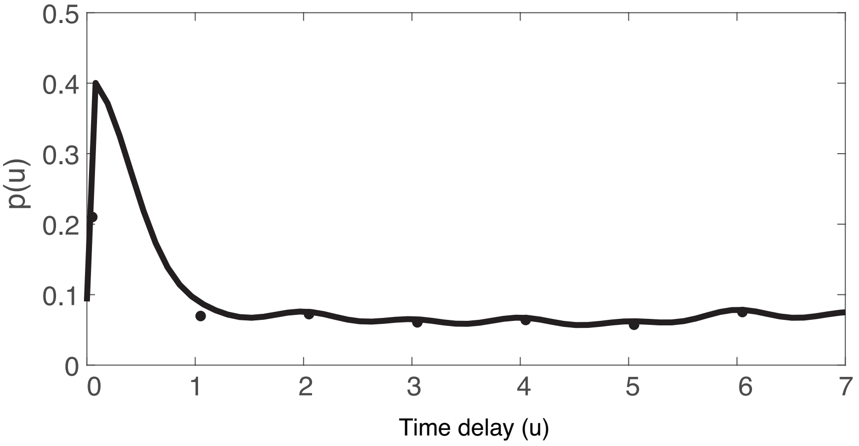


Figure S9:
